# Optimizing CRE and PhiC31 mediated recombination in *Aedes aegypti*

**DOI:** 10.1101/2023.07.07.548128

**Authors:** Leonela Z Carabajal Paladino, Ray Wilson, Priscilla YL Tng, Vishaal Dhokiya, Elizabeth Keen, Piotr Cuber, Will Larner, Sara Rooney, Melanie Nicholls, Anastasia Uglow, Luke Williams, Michelle AE Anderson, Sanjay Basu, Philip T. Leftwich, Luke Alphey

**Affiliations:** Arthropod Genetics Group, The Pirbright Institute, Pirbright, UK; The Department of Biology, University of York, York, UK; Department of Vector Biology and Department of Tropical Disease Biology, Liverpool School of Tropical Medicine, Liverpool, UK; School of Natural and Environmental Sciences, Newcastle University, Newcastle upon Tyne, UK; Molecular Laboratories, Core Research Laboratories, Natural History Museum, London, UK; School of Live Sciences, University of Warwick, Coventry, UK; Forest Research, Farnham, UK; Animal and Plant Health Agency, UK; Molecular Biology Team, R&D Division, Oxitec, Oxford, UK; School of Biological Sciences, University of East Anglia, Norwich, UK

**Author notes:** Correspondence: Luke Alphey.

**Keywords:** integration, plasmid, cassette exchange, transgenesis, mosquito

## Abstract

Genetic manipulation of *Aedes aegypti* is key to developing a deeper understanding of this insects’ biology, vector-virus interactions and makes future genetic control strategies possible. Despite some advances, this process remains laborious and requires highly skilled researchers and specialist equipment. Here we present two improved methods for genetic manipulation in this species. Use of transgenic lines which express Cre recombinase allowed, by simple crossing schemes, germline or somatic recombination of transgenes, which could be utilized for numerous genetic manipulations. PhiC31 integrase based methods for site-specific integration of genetic elements was also improved, by developing a plasmid which expresses PhiC31 when injected into early embryos, eliminating the need to use costly and unstable mRNA as is the current standard.

## 1 Introduction

*Aedes aegypti* is the key mosquito vector of a range of important arboviruses including dengue virus, Zika virus, chikungunya virus and yellow fever virus (World Health Organisation, 2014). A better understanding of its life cycle, methods of population control, and host-virus interaction, among others, is therefore important. Some studies, e.g. the generation of genetic control methods such as engineered sterile males or gene drives, benefit from the use of genetic manipulation of the mosquito (Alphey, 2014).

Genetic transformation of *Ae. aegypti*, i.e., stable integration of a foreign sequence (transgene), was first reported by Coates et al. (1998), who used the *mariner* transposable element to insert a wild-type copy of the *Drosophila melanogaster cinnabar* gene into an *Ae. aegypti* strain mutant for the homologous gene *kynurenine hydroxylase*. Since then, there has been a plethora of reports of the successful generation of *Ae. aegypti* transgenic lines for basic studies and directed towards developing functional strains for disease control. Despite significant improvements, the generation of transgenic mosquitoes is still laborious and a specialist activity available in relatively few labs. For this reason, alternative methods for transformation, and means to maximise the value and utility of each transgenic line, would be highly desirable.

*In vivo* recombination, primarily using the Cre-*lox* system, has been extremely valuable in animal genetics (Nagy, 2000). Initial attempts to use this tool in *Ae. aegypti* indicated a rather low efficiency (Jasinskiene et al., 2003). Perhaps as a result, the system has been subsequently under-utilised, with the exception of some “cassette exchange” transformation experiments (Nimmo et al., 2006; Haghighat-Khah et al., 2015; Häcker et al., 2017).

Cre recombinase is an enzyme expressed by the coliphage P1 that recognises short asymmetrical *lox* sequences and catalyses their recombination (Austin et al., 1981; Sauer, 1987). When both *lox* sequences have the same orientation, the recombination event will result in the deletion of the fragment between both *lox* sites, leaving only one *lox* sequence behind (Figure 1a).

**Figure 1:**
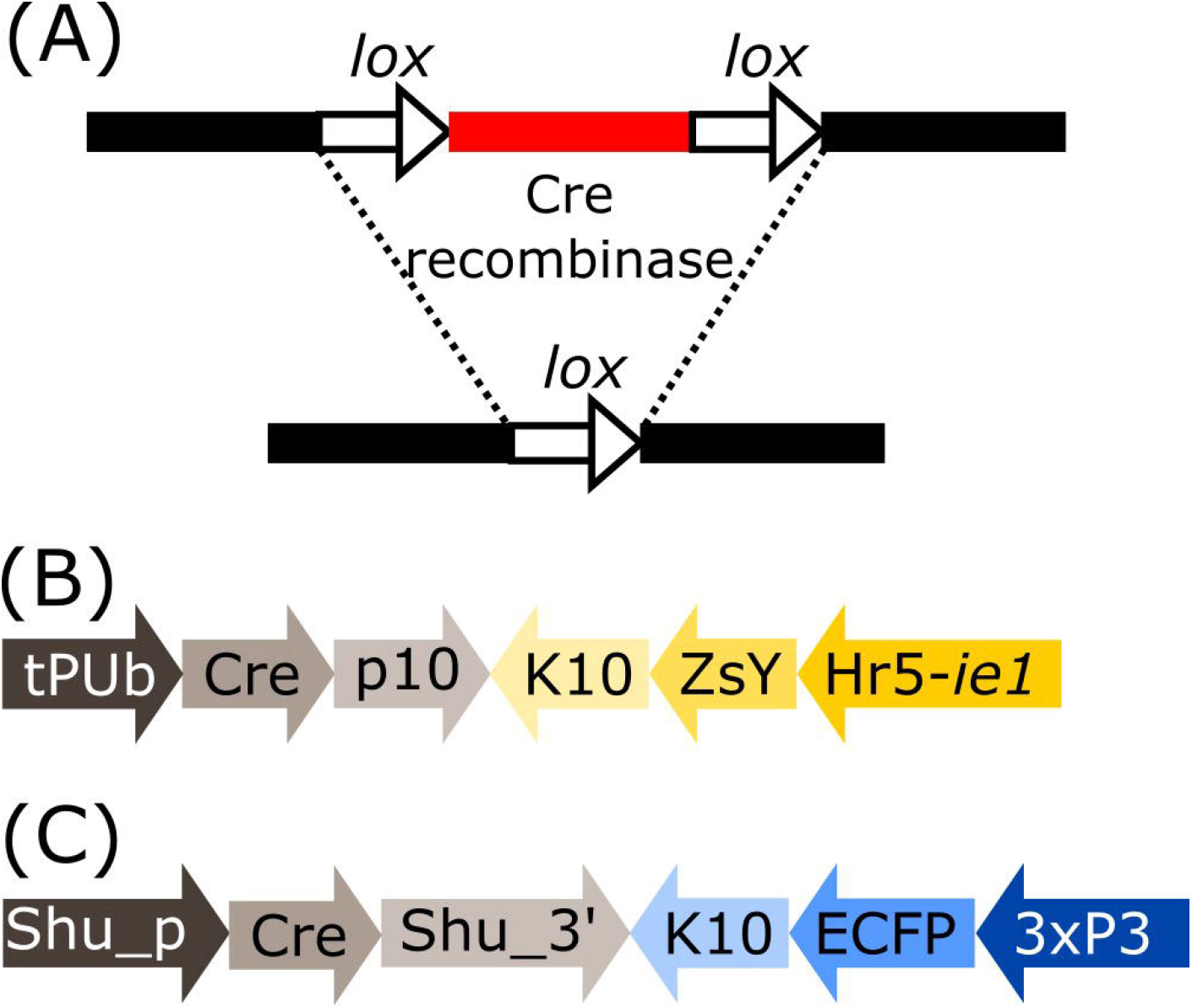
**(A)** Schematic representation of the removal of a DNA fragment (red) leaving one *lox* sequence behind (white arrow). **(B)** Schematic representation of plasmid tPUb-Cre which includes the elements necessary for the ubiquitous expression of Cre recombinase (grey segments) and a transformation marker providing yellow fluorescence in most tissues (yellow segments: yellow body transformation marker, Hr5-*ie1*-ZsYellow). **(C)** Schematic representation of plasmid Shu-Cre which includes the elements necessary for germline expression of Cre recombinase (grey segments) and a blue eye-specific transformation marker (blue segments: 3xP3-ECFP). tPUb: truncated *Polyubiquitin* promoter, Cre: Cre recombinase nucleotide sequence, p10: *Autographa californica* nucleopolyhedrovirus (AcMNPV) *p10* 3’UTR, K10: *Drosophila melanogaster fs(1)K10* 3’UTR, ZsY: *Zoanthus* sp. yellow fluorescent protein, Hr5-*ie1*: AcMNPV *ie1* promoter fused with homologous region 5 (hr5) enhancer, Shu_p: s*hut-down* promoter, Shu_3’: *shut-down* 3’UTR, ECFP: enhanced cyan fluorescent protein, 3xP3: three tandem repeats of Pax-6 homodimer binding site fused to a basal promoter element. Cre-expressing plasmids have additional elements not shown, e.g., *piggyBac* terminal sequences.

The native *lox*P sequence from phage P1 comprises two inverted repeats separated by a spacer region (Hoess et al., 1982). The efficiency of the recombination is determined by the homology within the spacer region (Hoess et al., 1986), while the affinity of the recombinase is determined by the inverted repeat palindromic arms (Albert et al., 1995). Genome screening and directed mutation of the spacer region have generated several additional *lox* sites that can be used for genome manipulation (Albert et al., 1995; Araki et al., 1997; Lee and Saito, 1998; Langer et al., 2002; Livet et al., 2007).

The use of Cre recombinase was previously described in *Ae. aegypti*, based on micro-injection of either purified Cre enzyme, capped mRNA helper, or a plasmid encoding Cre (Jasinskiene et al., 2003; Nimmo et al., 2006; Haghighat-Khah et al., 2015; Häcker et al., 2017). Here we revisited the feasibility of *in vivo* recombination using Cre expressed both in germline to produce offspring with modified genomes, and in somatic cells to produce somatic mosaics. We tested the efficiency of the transgenic mosquito lines expressing Cre, in different constructs that can then be modified for a variety of studies, and we were able to achieve each intended manipulation with good efficiency.

Another useful recombinase, which may be used in combination with Cre-*lox*, is the integrase enzyme expressed by *Streptomyces* temperate phage PhiC31, that catalyses the unidirectional and stable integration of large DNA fragments in specific sites. In synthetic biology applications, the enzyme catalyses the recombination of the phage attachment site (*attP*) typically inserted into the target genome after using conventional transgenesis, and the bacterial attachment site (*attB*) present in the donor plasmid. As a result, new hybrid *attL* and *attR* sites are generated. Unlike *lox* sites, *attP* and *attB* are substantially different in sequence, so the recombinant sequences *attL* and *attR* do not match either and are no longer recognised by the integrase (Thorpe and Smith, 1998). Consequently, recombination mediated by PhiC31 integrase is essentially irreversible. This has allowed its use for germline transformation, whereas with Cre-*lox* recombination the equilibrium of the reversible reaction is directed heavily towards the non-integrated form. Though mutant derivatives of *lox* can potentially be used to reduce the rate of excision, initial experiments indicated PhiC31 to be much preferable (Nimmo et al., 2006).

Transformation of *Ae. aegypti* using PhiC31 integrase is traditionally performed by expressing the enzyme from mRNA included in the injection mix (Nimmo et al., 2006; Franz et al., 2011; Khoo et al., 2013; Haghighat-Khah et al., 2015). The use of mRNA adds extra steps to the transformation procedure, as it needs to be transcribed *in vitro*, DNase treated, purified, precipitated, and resuspended. Efficiency, convenience and reproducibility are also potentially affected by the relative instability of this mRNA. Here we improved this tool by expressing the PhiC31 enzyme from a plasmid included in the injection mix, avoiding the use of mRNA.

## 2 Materials and Methods

### 2.1 Plasmid construction

Plasmids were generated by standard molecular biology techniques including HiFi assembly (New England BioLabs). All plasmids were prepared for microinjection using the NucleoBond Xtra Midiprep kit EF (Machery-Nagel) and confirmed by Sanger sequencing. Sequences of the plasmids designed for this manuscript and of the transgenes of the transgenic lines used are deposited into GenBank, and the accession numbers are detailed in Supplementary Table 1.

### 2.2 General mosquito maintenance

*Aedes aegypti* mosquitoes (Liverpool wild-type strain and transgenics derived from them) were reared at 28 °C and 70% relative humidity with a 14:10 h light:dark cycle. Adults were kept in BugDorm cages with cotton wool pads soaked with 10% sucrose solution. Transgenic lines were maintained in hemizygous state by crossing transgenic males with wild-type females at a 1:2 male:female ratio. From day 5-7 post-eclosure, females were blood fed once every five days, three times, to collect three ovipositions. Defibrinated horse blood (TCS) was offered until most of the females were engorged using a Hemotek membrane feeding system (Hemotek Ltd) covered with hog gut and an over-layer of parafilm. Two days after blood meal, egg cups consisting of 1 oz clear polypropylene portion pots with a strip of coffee filter paper wrapped inside and half filled with reverse osmosis water, were added to the cages to allow egg laying. The egg cups were removed 5 days later, and egg papers hatched when required. Eggs were vacuum hatched for 2 hours in 50 ml solution of reverse osmosis water with LiquiFry N° 1 (Interpet) (500 ml of water + 6 drops of LiquiFry). The larvae were then transferred to a new tray with reverse osmosis water and fed TetraMin ornamental Fish Flakes *ad libitum*. Transgenic mosquitoes were identified in larval or pupal stages based on their fluorescent marker profile using an MZ10F stereomicroscope (Leica Microsystems) equipped with suitable filters. Mosquitoes were sexed under a stereomicroscope at the pupal stage according to the shape of the genital lobe.

### 2.3 Generation of transgenic lines

Cages with approximately 1,000 5-7 day post-eclosure wild-type females mated with 500 wild-type males were blood fed as described above in the general mosquito maintenance section. Five to seven days after the blood meal, 15-20 females were transferred into a *Drosophila* tube with damp cotton wool covered by a layer of filter paper and kept for 20-30 mins in the dark to promote oviposition. White eggs were lined up side by side in the same orientation on a piece of filter paper and then transferred onto a cover slip with Scotch Double Sided Tape 665 (3M). Eggs were covered with halocarbon oil 27 (Sigma Aldrich) to prevent desiccation before microinjection in their posterior end. Injections were performed using Sutter quartz needles (1.0 mm external diameter 0.70mm internal diameter needles with filaments) drawn out on a Sutter P-2000 laser micropipette puller (Sutter Instrument) with the following program: Heat = 729, FIL = 4, VEL = 40, DEL = 128, PUL = 134, LINE = 1. Injections were carried out using a standard microinjection station equipped with a FemtoJet 4x microinjector (Eppendorf). The injection mix contained 500 ng/μl of the plasmid intended for insertion, 300 ng/μl of helper plasmid AePUb-hyperactive *piggyBac* transposase (Anderson et al. 2022) for the generation of transgenic lines or AePUb-PhiC31 (this manuscript) for PhiC31 mediated integration, 1x Injection buffer (Coates et al., 1998) and endotoxin-free water.

After microinjection, the eggs were washed with water and transferred onto damp filter paper in a moist chamber. Five days after injection the eggs were vacuum hatched as described for general maintenance. The collected pupae were sexed under a stereomicroscope. Each Gen_0_ male was single crossed with 5 wild-type females, and the individuals were pooled in groups of 3-5 single crosses after 48 hours. Gen_0_ females were pooled in groups of 4-5 females and 8-10 wild-type males. The cages were fed with 10% sucrose, blood fed every 5-6 days and 5 ovipositions were collected in total. The Gen_1_ egg papers were hatched as previously described, and the 3rd-4th instar larvae were screened for the presence of the transformation marker. Single positive Gen_1_ individuals were crossed with wild-type counterparts and maintained as described above.

The number of transgene insertions and sex-linkage of the constructs were assessed according to standard rearing procedures by crossing transgenic males to wild-type females and quantifying the number of males and females of wild-type and transgenic phenotypes, and comparing them to the expected frequencies for an autosomal single insertion (i.e. 50% males and 50% females, 50% transgenic and 50% wild-type). Flanking PCR was also used (see protocol in the Molecular analysis section). Only lines with single insertions were used in experiments. Information regarding the insertion site of the lines used in these experiments is detailed in Supplementary Table 2.

### 2.4 Experimental crosses

50 females and 25/50 males of the corresponding mosquito lines (wild-type or transgenics) were pooled together according to the experiment (see Results section) and maintained as described above. For screening, 3 replicates of 300 eggs each were hatched and kept in trays with 1.5 l of reverse osmosis water and fed with TetraMin ornamental Fish Flakes *ad libitum*. This was done to ensure optimal developmental conditions for all the genotypes. Individuals were screened in larval or pupae stage and photographed when needed using a DFC7000 T camera (Leica Microsystems).

### 2.6 Statistical analysis

Chi square tests of independence were used in crosses involving the tPUb-Cre and Shu-Cre lines to study the egg to pupa survival of the double hemizygous, both single hemizygous and wild-type individuals of the different crosses. The observed values were calculated as an average of those observed in the three replicates. For individuals carrying a construct susceptible to recombination, the values of the non-recombined and recombined phenotypes were added. The expected values were calculated considering a normal segregation of the constructs and full survival of the phenotype, i.e., 25% of the average of the total number of individuals recovered per replicate.

Student’s *t*-tests were carried out to compare number of individuals per genotype between crosses of the Shu-Cre line.

All tests were carried out using R (ver 4.2.2) (R Core Team, 2022) in RStudio 2022.12.0+353, with the MASS (Venables and Ripley, 2002), rstatix (Kassambara, 2023) and dplyr (Wickham et al., 2023)

### 2.6 Molecular analysis

Genomic DNA was extracted from larvae or pupae of the selected lines using NucleoSpin Tissue gDNA extraction kit (Macherey-Nagel).

Flanking PCR was performed according to previously reported methods (Liu and Chen, 2007; Martins et al., 2012). Genomic DNA was digested using enzymes DpnII, MspI and NcoI (New England Biolabs) and PCRs were performed with DreamTaq (Thermo Fisher Scientific). Adaptors were built with primers 6770 (5’-GTGTAGCGTGAAGACGACAGAAAGGGCGTGGTGCGGAGGGCGGTG-3’) and 6776 (5’-GATCCACCGCCCTCCG-3’) for DpnII, 6770 and 6775 (5’-CGCACCGCCCTCCG-3’) for MspI, and 6770 and 6789 (5’-CATGCACCGCCCTCCG-3’) for NcoI. The primary PCR used primers 6774 (5’-GTGTAGCGTGAAGACGACAGAA-3’) and 6772 (5’-CAGTGACACTTACCGCATTGACAAG-3’) to amplify the 5’ insertion site, and 6774 and 6773 (5’-CAGACCGATAAAACACATGCGTCA-3’) to amplify the 3’ insertion site. The nested PCR used primers 6774 and 6771 (5’-GGCGACTGAGATGTCCTAAATGCAC-3’) to amplify the 5’ insertion site, and 6774 and 6787 (5’-ACGCATGATTATCTTTAACGTACG-3’) to amplify the 3’insertion site.

PCR reactions for the other experiments were performed in a final volume of 100 μl using 50-150 ng of template, 5 μl of primer forward 10 μM, 5 μl of primer reverse 10 μM, 50 μl of Q5 High-Fidelity 2x Master Mix (New England Biolabs) and water.

To test for the removal of internal sequences for interaction analyses, primers 6676 (5’-TGTGCAGTCGGTTAGTTGGGAAAGG-3’) and 5038 (5’-GGCCATTGTGACTTTGAAGGTGGAGG-3’) were used. Cycling conditions included an initial denaturation step at 98 degrees for 30 s, 35 cycles of 98 degrees 10 s, 72 degrees 1 min 35 s, and a final elongation at 72 degrees for 2min. A fragment of 3135 bp was expected if the internal sequence was present, and of 350 bp if the sequence was absent.

To test for the removal of backbone sequences, primers 5842 (5’-CAGACATGATAAGATACATTGATG-3’) and 6792 (5’-AATGACATCATCCACTGATCG-3’) were used. Cycling conditions included an initial denaturation step at 98 degrees for 30 s, 35 cycles of 98 degrees 10 s, 59 degrees 15 s and 72 degrees 2 min 45 s, and a final elongation at 72 degrees for 2min. A fragment of 5281 bp was expected if the backbone sequence was present, and of 578 bp if the sequence was absent.

To test for correct PhiC31 integrase mediated integration, primer pair 6707 (5’-TCGGTCTGTATATCGAGGTTTAT-3’) and 6799 (5’-CCCTTCACGGTGAAGTAGTG-3’) was used to amplify the *attL* region, and primer pair 6795 (5’-GCACAAGCTGGAGTACAACTA-3’) and 6791 (5’-CCAGTTCGGTTATGAGCCGT-3’) was used to amplify the *attR* region. Cycling conditions included an initial denaturation step at 98 degrees for 30 s, 35 cycles of 98 degrees 10 s, 63 degrees 15 s and 72 degrees 45 s, and a final elongation at 72 degrees for 2min. Fragments of 1101 bp and 1393 bp were expected for *attL* and *attR*, respectively.

PCR products were run in a 1% agarose gel with SYBR™ Safe DNA Gel Stain (Thermo Fisher Scientific) and photographed using a Bio-Rad Gel Doc imaging system. As clear single bands were observed in all cases, the remaining PCR product was purified using NucleoSpin Gel and PCR Clean-up kit (Macherey-Nagel) and Sanger sequenced using both forward and reverse primers.

## 3 Results

### 3.1 Cre recombinase expressing lines

We generated transgenic mosquito lines expressing Cre recombinase either ubiquitously using a truncation of the *Polyubiquitin* gene promoter (AAEL003877, Bartholomeeusen *et al*., 2018) (tPUb-Cre, Figure 1b) or the germline specific *shut-down* gene promoter (from OP823146, Anderson *et al*., 2023) (Shu-Cre, Figure 1c). The Cre coding sequence used was modified for expression in *Ae. aegypti* by adjusting codon usage to avoid rare codons (“codon-optimised”). We tested these lines by crossing them to different transgenic reporter lines.

### 3.2 Somatic effect of Cre recombinase expression

To analyse these lines for their somatic effect, we generated a reporter mosquito line carrying the construct PUb-loxP-mCh-stop-loxP-AmC (Figure 2a). These individuals express the red fluorophore mCherry over the whole body; after Cre mediated recombination, the mCherry open reading frame should be removed and the cyan fluorophore AmCyan expressed instead (Figure 2a, Supplementary Figure 1).

**Figure 2:**
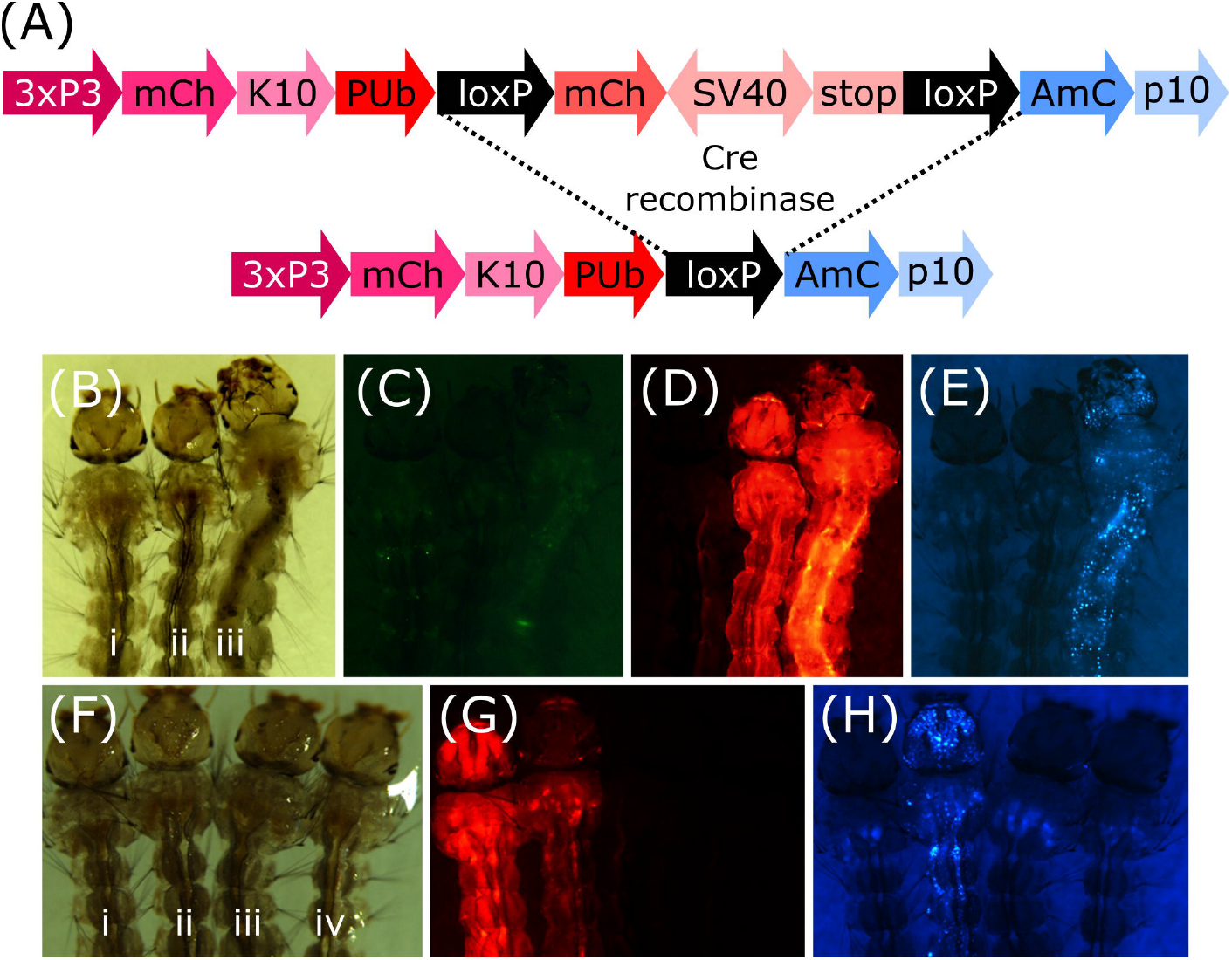
**(A)** Schematic representation of plasmid PUb-loxP-mCh-stop-loxP-AmC before and after Cre mediated recombination. **(B-E)** Individuals hemizygous for tPUb-Cre (i) or PUb-loxP-mCh-stop-loxP-AmC (ii) and double hemizygous (iii). **(F-H)** Individuals hemizygous for PUb-loxP-mCh-stop-loxP-AmC (i) or Shu-Cre (iii), double hemizygous (ii) and wild-type (iv). **(B)** and **(F)**: bright field, **(C)**: ZsYellow filter, **(D)** and **(G)**: mCherry filter, **(E)** and **(H)**: AmCyan filter. 3xP3: three tandem repeats of Pax-6 homodimer binding site fused to a basal promoter element, mCh: mCherry red fluorescent protein, K10: *Drosophila melanogaster K10* 3’UTR, PUb: *Polyubiquitin* promoter, SV40: Simian virus 40 PolyA sequence, AmC: *Anemonia majano* cyan fluorescent protein, p10: *Autographa californica* nucleopolyhedrovirus *p10* 3’UTR.

We screened the F_1_ progeny of crosses between hemizygous tPUb-Cre or Shu-Cre individuals with hemizygous counterparts from the reporter line. The three transgenic lines have their own transformation marker making it possible to identify all the genotypes of the F_1_: yellow body for tPUb-Cre, blue optic nerves for Shu-Cre, and red optic nerves for the reporter (Figures 1a-b and 2a).

For both types of crosses, all double hemizygous F_1_ individuals expressed mCherry and AmCyan over the whole body (Figure 2b-h), indicating that the removal of the loxP-mCh-stop-loxP section (leaving one *lox*P sequence behind) occurred in most but not all cells. Whole body expression of Cre in the tPUb-Cre line was expected due to the nature of the promoter. In the Shu-Cre line, although expression of *shut-down* is germline-specific, the promoter fragment used was known to provide some somatic expression as well (Anderson et al., 2023), which was now visually corroborated.

### 3.3 Germline effect of Cre recombinase expression

To analyse the lines for their germline effect we generated a reporter mosquito line carrying the construct loxN-R-loxP-loxN-Y-loxP (Figure 3, Supplementary Figure 2). These mosquitoes express a red body marker from fragment R, and a yellow optic nerves marker from fragment Y. As lines tPUb-Cre and Shu-Cre have yellow body and blue optic nerves markers respectively, all the genotypes deriving from the crosses could be identified (see example in Supplementary Figure 3). The reporter line includes other sequences that are not relevant for this study (Supplementary Figure 2), and hence their effect is not addressed. The use of two overlapping pairs of incompatible *lox* sites – *lox*P and *lox*N – means that recombination between one pair removes one of the other pair, thus preventing recombination based on that site. Recombination therefore has two possible stable outcomes (i) recombination between *lox*P sites deleting segment Y, retaining segment R and therefore red fluorescence; (ii) recombination between *lox*N sites deleting segment R, retaining segment Y and therefore yellow fluorescence.

**Figure 3:**
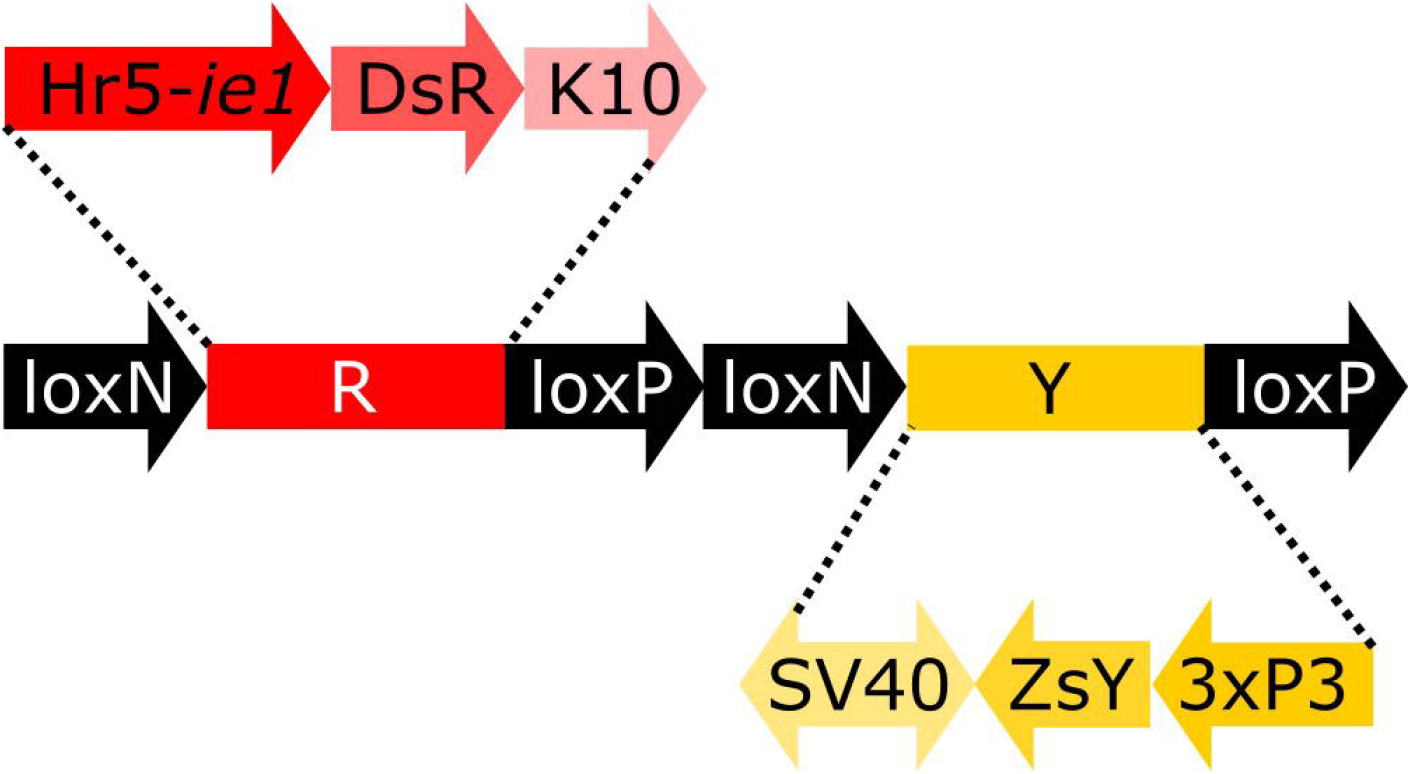
Schematic representation of construct loxN-R-loxP-loxN-Y-loxP. Hr5-*ie1*: AcMNPV *ie1* promoter fused with homologous region 5 (hr5) enhancer, DsR: *Discosoma* sp. red fluorescent protein, K10: *Drosophila melanogaster K10* 3’UTR, SV40: Simian virus 40 PolyA sequence, ZsY: *Zoanthus* sp. yellow fluorescent protein, 3xP3: three tandem repeats of Pax-6 homodimer binding site fused to a basal promoter element.

We first screened the F_1_ progeny of crosses between tPUb-Cre or Shu-Cre with the reporter line. In order to address any possible sex specific effect, both reciprocal crosses were performed for line tPUb-Cre, but as line Shu-Cre is linked to the *m* allele of the *M*/*m* male determining locus (“*m*-linked”), only Shu-Cre females were used. Double hemizygous F_1_ males and females were subsequently crossed to wild-type counterparts, the F_2_ was screened, and their phenotypes were assessed for recombination events.

The results for line tPUb-Cre are summarised in Figure 4a and detailed in Supplementary Tables 3 to 6. The results for line Shu-Cre are summarised in Figure 4b and detailed in Supplementary Tables 7 and 8. The analysis of the segregation of the tPUb-Cre and loxN-R-loxP-loxN-Y-loxP constructs (adding up recombined and non-recombined versions of the latter), showed no statistically significant differences with the expected frequencies (25% each - double hemizygous, single hemizygous for each insertion and wild-type) (Chi-square test of independence: p > 0.05). This suggests no substantial lethal effect of the recombinase line or the recombined fragments (power analysis: N > 250, β > 0.9, ω < 0.3 for all crosses). Crosses with the tPUb-Cre line produced only one recombinant individual for one of the two possible phenotypes (red body from red body or yellow optic nerves). This indicated little or no expression – or function, but see below – of Cre in the germline of this strain, likely due to the PUb promoter fragment used in this line.

**Figure 4:**
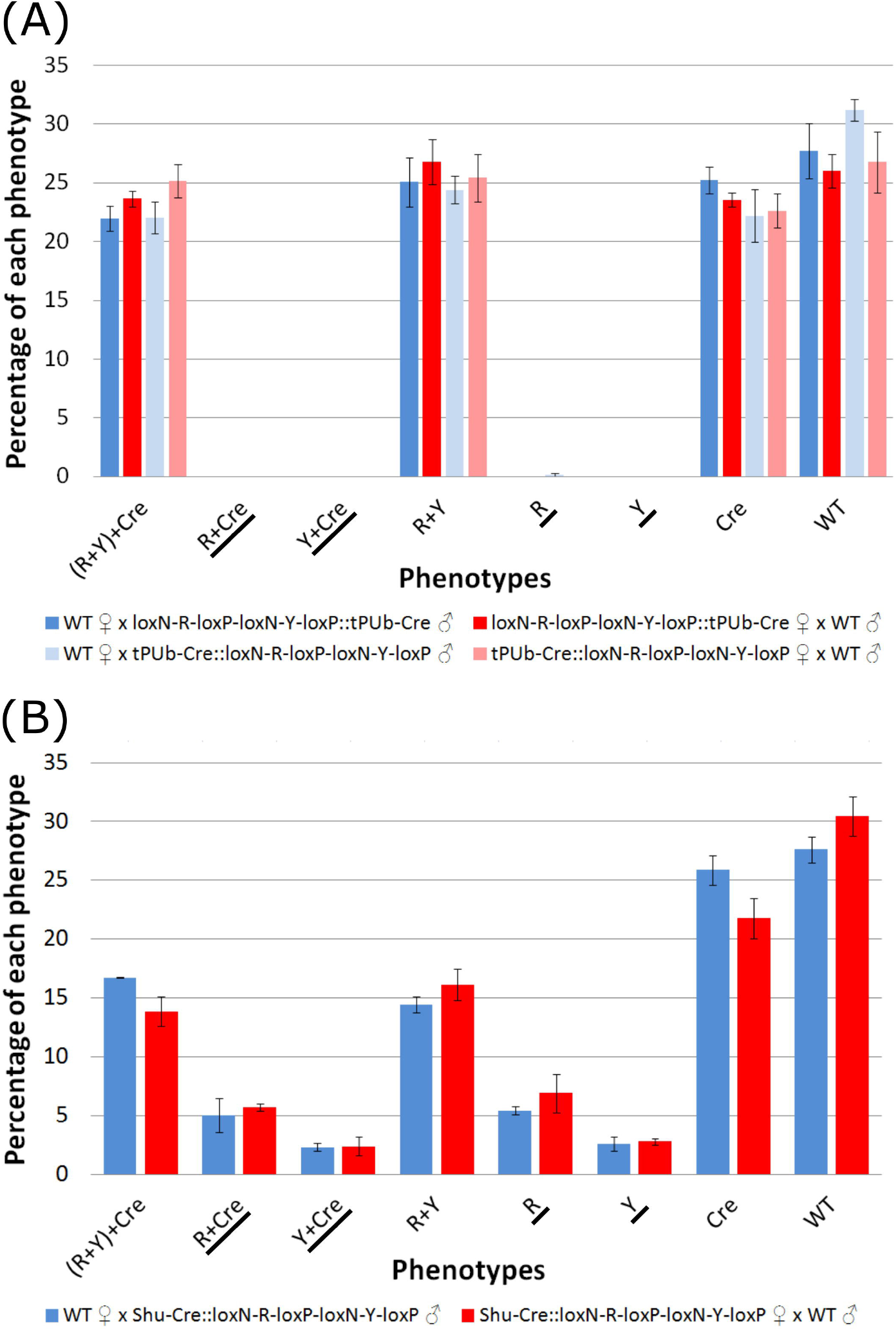
**(A)** Phenotypes of F_2_ individuals from crosses between double hemizygous loxN-R-loxP-loxN-Y-loxP and tPUb-Cre with wild-type counterparts. **(B)** Phenotypes of F_2_ individuals from crosses between double hemizygous loxN-R-loxP-loxN-Y-loxP and Shu-Cre with wild-type counterparts. Bars represent the average obtained from the three trays analysed, and the error bars are the standard error. Recombined phenotypes are underlined. R+Y: loxN-R-loxP-loxN-Y-loxP, R: fragment without segment Y= loxN-R-loxP, Cre: tPUb-Cre in A and Shu-Cre in B, Y: fragment without segment R= loxN-Y-loxP, WT: wild-type.

Crosses with the Shu-Cre line generated individuals of each of the two possible recombined phenotypes. Analysis of the observed frequencies of the segregating constructs found no evidence for lethal effect for any of the genotypes (Chi-square test of independence: p > 0.05), and expression of Cre recombinase in the germline. A linear model was constructed to compare the number of individuals per phenotype between crosses, and no differences were observed (Student’s *t*-test: p > 0.05, Supplementary Table 9). A power analysis indicates that there is no evidence for a difference in efficiency of the Cre recombinase between the male and female germline (β = 0.9, *f* = 0.2).

The efficiency of the Cre lines in the germline, calculated as the percentage of individuals carrying a recombined construct (R+Cre, Y+Cre, R and Y) was 0.16% for line tPUb-Cre and between 15.4 and 17.8% for the Shu-Cre line.

### 3.4 Further experiments using Shu-Cre

As the Shu-Cre line was found to be highly efficient at inducing recombination in the germline (15.4-17.8% efficiency), we further analysed it using other lines containing *lox* sites.

#### 3.4.1 Removal of internal sequences for interaction analyses

We used the Shu-Cre line to remove an internal fragment within a transgene of mosquito lines available in the laboratory. These lines were developed to help compare bipartite expression using the tet-off system (Gossen and Bujard, 1992) (promoter-tTA>TRE-reporter) with direct expression (promoter-reporter). Here we focus on the Cre-*lox*N data only.

We used two lines carrying the reporter construct A-loxN-B-loxN-C (Figure 5) in two different locations on chromosome 2: lines D and E (Supplementary Table 2).

**Figure 5:**
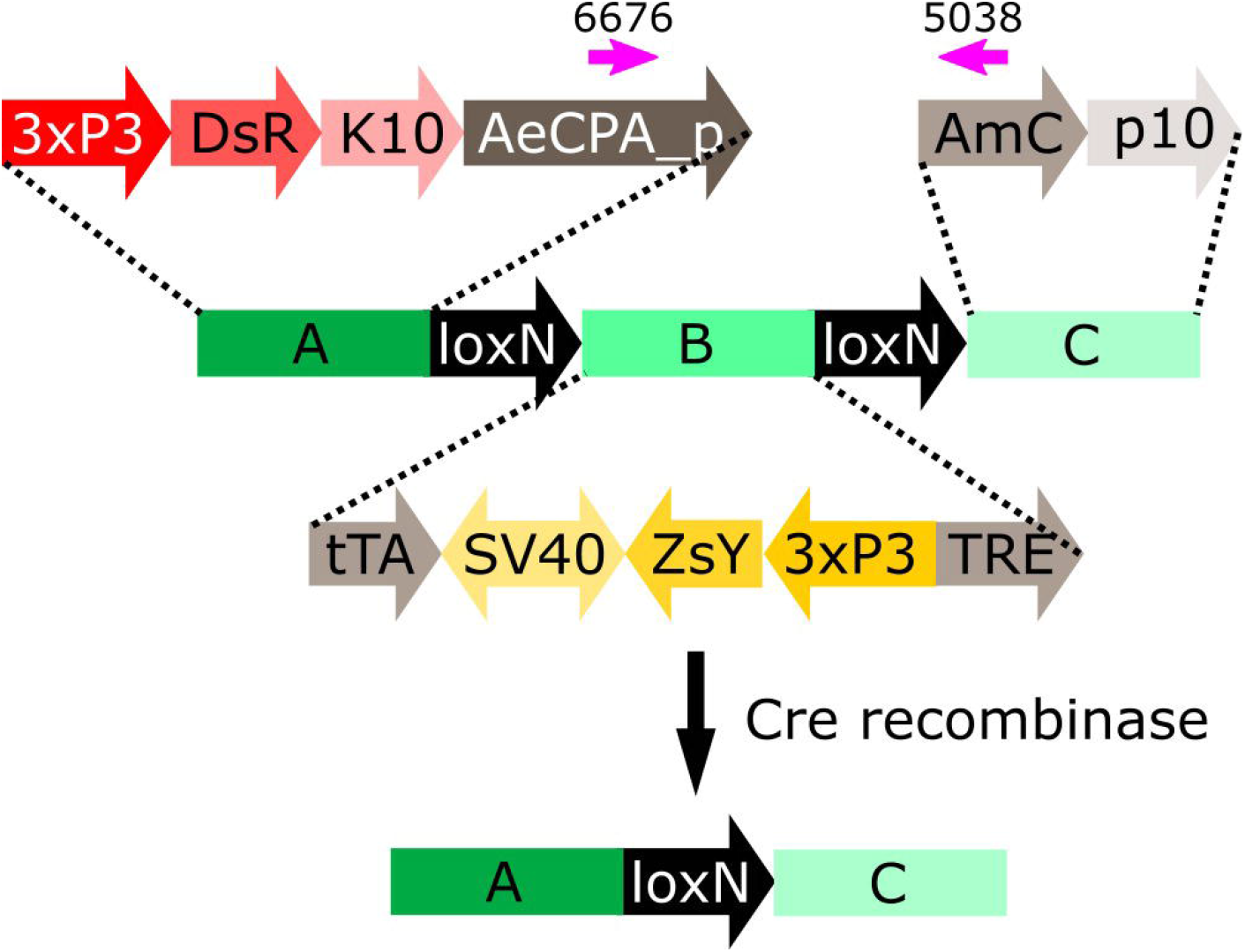
Schematic representation of construct A-loxN-B-loxN-C. The pink arrows indicate the location of the primers used for validation of the removal of fragment B. 3xP3: three tandem repeats of Pax-6 homodimer binding site fused to a basal promoter element, DsR: *Discosoma* sp. red fluorescent protein, K10: *Drosophila melanogaster K10* 3’UTR, AeCPA_p: *Aedes aegypti Carboxypeptidase A* promoter, tTA: tetracycline-controlled transactivator, SV40: Simian virus 40 PolyA sequence, ZsY: *Zoanthus* sp. yellow fluorescent protein, TRE: tetracycline response element, AmC: *Anemonia majano* cyan fluorescent protein, p10: *Autographa californica* nucleopolyhedrovirus *p10* 3’UTR.

The construct A-loxN-B-loxN-C has two transformation markers: red optic nerves for segment A, and yellow optic nerves for segment B (Figure 5). The removal of fragment loxN-B-loxN (leaving a *lox*N behind) should result in the loss of the yellow marker (Figure 5).

Males of each line carrying the construct A-loxN-B-loxN-C were crossed to Shu-Cre females, and the double hemizygous F_1_ were crossed to wild-type counterparts. The F_2_ was screened; results are summarised in Figure 6a and detailed in Supplementary Tables 10 and 11. Analysis of the segregation of the constructs Shu-Cre and A-loxN-B-loxN-C (adding up non-recombined A-loxN-B-loxN-C and recombined A-loxN-C individuals) showed no significant statistical differences (p > 0.05) with the expected values (25% each - double hemizygous, single hemizygous for each insertion and wild-type). It is unclear, however, why fewer A-loxN-B-loxN-C individuals were observed compared to double hemizygous Shu-Cre::A-loxN-B-loxN-C, as no effect on survival is expected of the Shu-Cre construct (as observed when the germline effect of the line was tested above).

**Figure 6:**
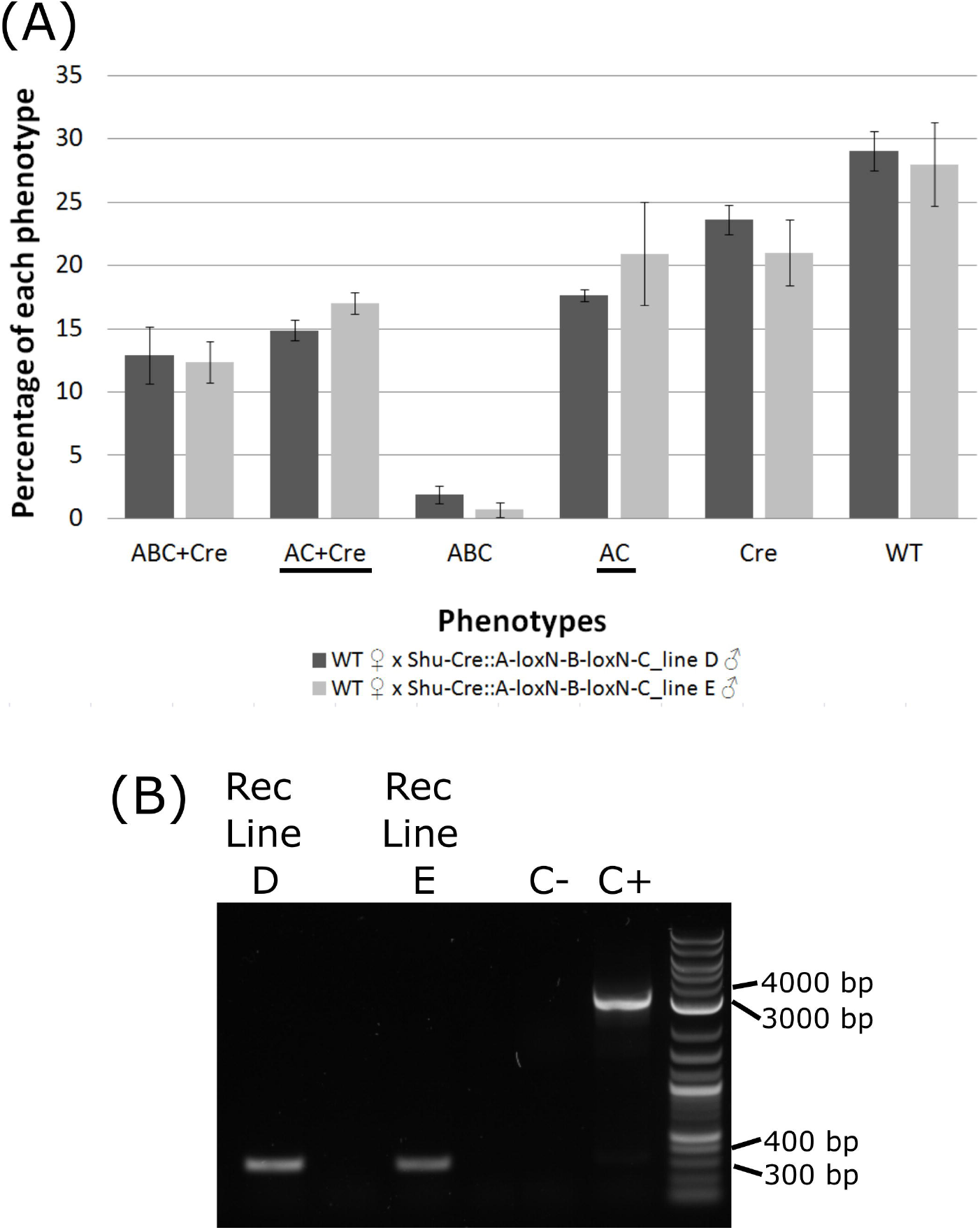
**(A)** Genotypes of F_2_ individuals from crosses between double hemizygous A-loxN-B-loxN-C and Shu-Cre with wild-type counterparts. Bars represent the average obtained from the three sets of progeny analysed, and the error bars are the standard error. Recombined phenotypes are underlined. **(B)** PCR validation of the removal of fragment B. The pool of recombinant individuals from lines D and E show an amplicon of between 300 and 400 bp, expected after the removal of fragment B. The positive control (C+) using purified transformation plasmid as template generated an amplicon of between 3000 and 4000 bp, expected if fragment B is present. ABC+Cre: double hemizygous carrying A-loxN-B-loxN-C and Shu-Cre, AC+Cre: double hemizygous carrying the recombined fragment A-loxN-C (B fragment removed) and Shu-Cre, ABC: individuals carrying the A-loxN-B-loxN-C construct, Cre: individuals carrying Shu-Cre, AC: individuals carrying the recombined version A-loxN-C, WT: wild-type, Rec: recombined, C-: negative control (water).

The efficiency of the Cre line in the germline, calculated as the percentage of individuals carrying a recombined construct (Shu-Cre::A-loxN-C and A-loxN-C), was higher in this experiment with 32.49% for Line D and 37.93% for Line E (compared to 15.4-17.8% obtained with the loxN-R-loxP-loxN-Y-loxP lines – see previous section).

For a subset of the recombinants, the removal of fragment B was additionally assessed by PCR (Figure 6b). The PCR primers were located within sections A and C (Figure 5, pink arrows). Presence of fragment B should be detected by the amplification of an amplicon of 3135 bp, and the absence of the fragment should generate an amplicon of 350 bp. As observed in Figure 6b, recombined lines D and E showed an amplicon of the expected size, the negative control of water (C-) did not produce an amplicon, and the positive control using purified plasmid (C+) presented only the larger amplicon. The obtained fragments were purified and the identity of the sequences was confirmed *via* Sanger sequencing (Supplementary Table 12).

#### 3.4.2 Removal of backbone sequences

Methods that integrate an entire plasmid, e.g. PhiC31 mediated integration, insert in the target site sequences that may be undesirable for some downstream studies or uses, e.g. plasmid origin of replication and antibiotic resistance genes. With efficient Cre-*lox* recombination available as shown above, an option to remove undesirable sequences post-integration could be provided by flanking such sequences with suitable *lox* sites. We used the lines generated below using PhiC31 integrase (attL-TRE-AmC-Bb-attR-AeCPA-tTA, Figure 8d) and crossed males of each line to Shu-Cre females. We only used one pool per line since, as expected and validated (see PhiC31 experiment below), all the pools from the same line have the same insertion (Supplementary Table 13). We then selected F_1_ males with red body (marker of the target line) and blue eyes (marker of the inserted plasmid in the target line and the Shu-Cre line) (see Figure 8d for the schematic representation of the transgene in the target line, and Figure 1c for the representation of the Shu-Cre transgene). As both the target line and the Shu-Cre line share the blue eyes marker, hemizygous males from the target line were not readily distinguished from double hemizygous males. We used at least 50 males per line, crossed them to wild-type females, and screened their offspring.

Due to the design of the donor plasmid used (Figure 8c), the removal of the backbone after the insertion (Figure 8d) also results in the removal of the marker for the transgene (i.e., blue optic nerves, Figure 8e). An effective recombination will produce individuals carrying only the marker of the original *attP* target line (i.e., red body, see Figure 8b).

As it was not possible to accurately identify the phenotype of the parental male (i.e., to easily distinguish hemizygous from double hemizygous due to the shared blue optic nerves marker), no quantitative analysis was performed. For each line tested, progeny with markers indicating recombination (red body but not blue optic nerves) were readily recovered. One such individual was selected from each line and used for subsequent analysis by PCR and Sanger sequencing. The PCR primers were located within the SV40 segment before the backbone and the Hr5-*ie1* promoter of the marker after the backbone (Figure 8d and e, primers 5842 and 6792 - pink arrows). Presence of the backbone should be detected by the amplification of an amplicon of 5281 bp, and the absence of this region should generate an amplicon of 578 bp (see difference between Figure 8d and e). As observed in Figure 7, lines A, D and C generated an amplicon between 5000 and 6000 bp before the removal of the backbone (+Bb), and a band between 500 and 600 bp after recombination (-Bb), the negative control of water (C-) did not produce an amplicon. Sanger sequencing of the obtained amplicons demonstrated precise *lox*P-mediated recombination, and efficient removal of the backbone.

**Figure 7:**
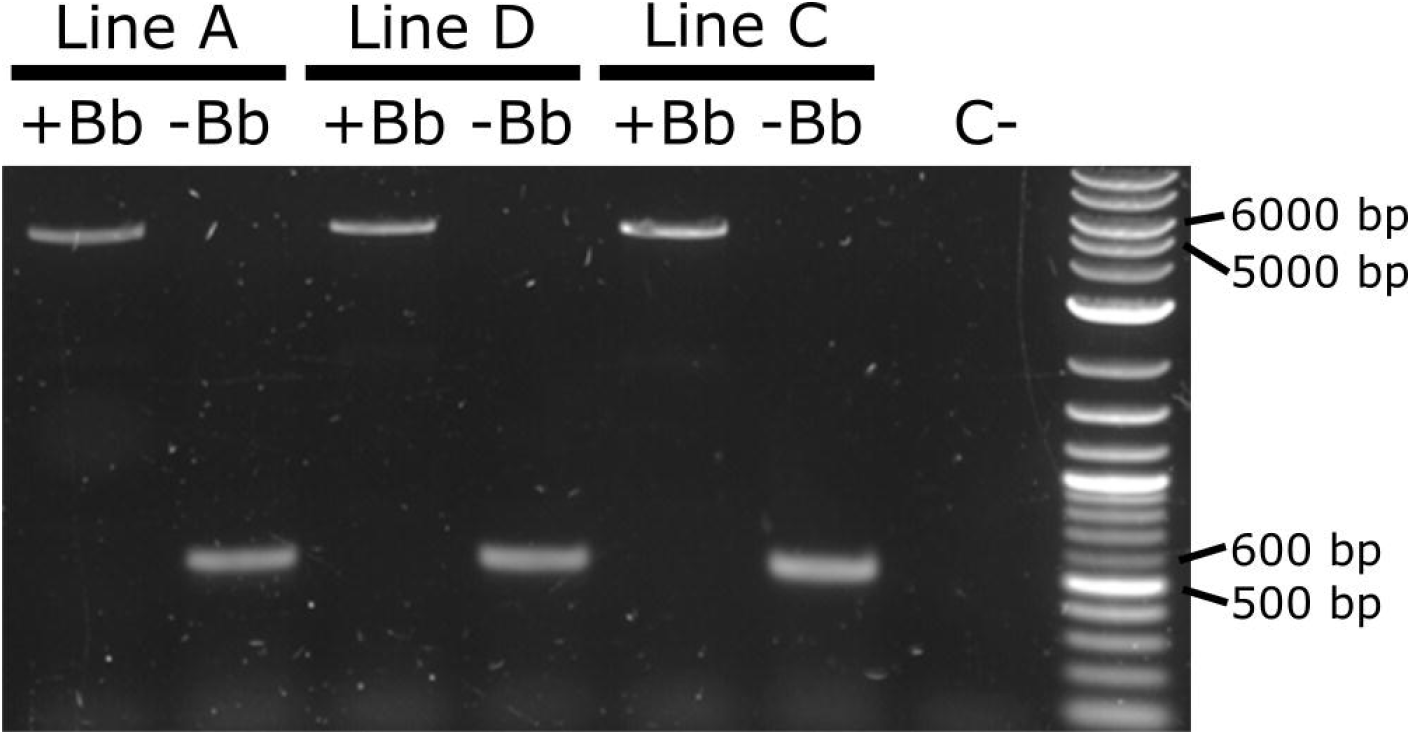
PCR amplification of the backbone (Bb) before and after crosses to Shu-Cre. Lines A, D and C were tested before (+Bb) and after (-Bb) the removal of the backbone of the inserted transgene, producing a large band between 5000 and 6000 bp when the backbone was present, and a small band between 500 and 600 bp once the backbone was removed. C-: negative control (water).

3.4.3 PhiC31 integrase mediated integration

Here we expressed the PhiC31 integrase (Genbank KT894025) in the injection mix from a plasmid using the *Ae. aegypti Polyubiquitin* promoter (Anderson et al., 2010) (Figure 8a). As target lines we used three mosquito lines available in the laboratory carrying a red body transformation marker (Figure 8b). In line A the transgene is in chromosome 1, in line D it is located in chromosome 2, and in line C it is located in chromosome 3 (Supplementary Table 2). As donor plasmid we designed the construct attB-TRE-AmC which contains a blue optic nerve marker (Figure 8c). An efficient integration of the donor plasmid will result in individuals carrying two markers, red body and blue optic nerves.

**Figure 8:**
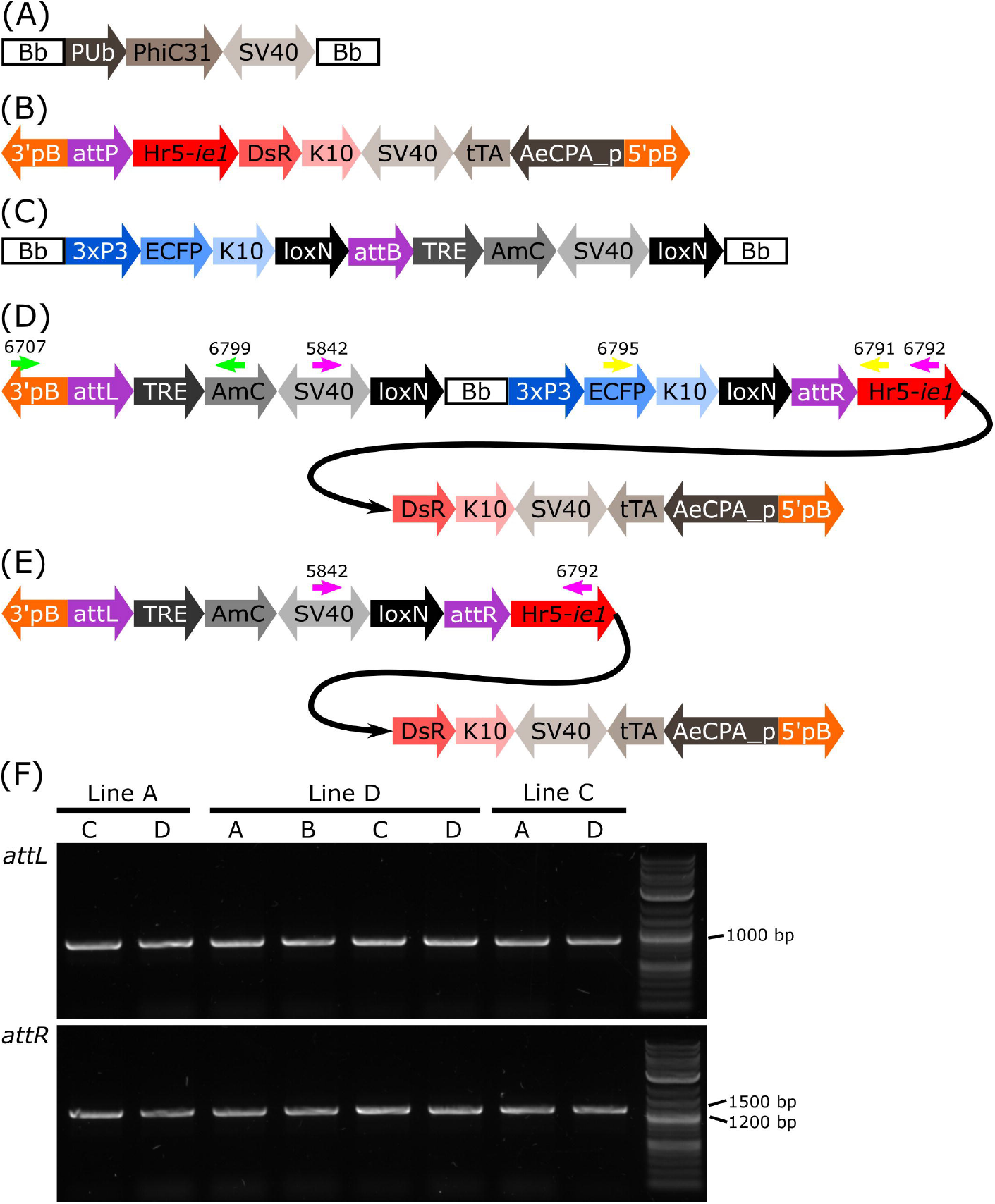
**(A)** Schematic representation of plasmid PUb-PhiC31. **(B)** Schematic representation of the target site. **(C)** Schematic representation of the donor plasmid. **(D)** Schematic representation of the donor plasmid inserted in the target site, and location of primers for PCR testing of *attL* and *attR* junctions and backbone removal (green, yellow and pink arrows, respectively). **(E)** Schematic representation of the insertion after removal of the backbone. **(F)** PCR amplification of junctions including sites *attL* and *attR* in all the pools with positive G_1_ (Table 1). All lines produced an amplicon of approx. 1000 bp for *attL*, and between 1200 and 1500 bp for *attR*, as expected. Bb: backbone, PUb: *Polyubiquitin* promoter, PhiC31: *Streptomyces* temperate phage PhiC31 integrase, SV40: Simian virus 40 PolyA sequence, 3’pb: 3’ piggyBac end, attP: phage attachment site, Hr5-*ie1*: *Autographa californica* nucleopolyhedrovirus *ie1* promoter fused with homologous region 5 (hr5) enhancer, *Discosoma* sp. red fluorescent protein, K10: *Drosophila melanogaster K10* 3’UTR, tTA: tetracycline-controlled transactivator, AeCPA_p: *Aedes aegypti Carboxypeptidase A* promoter, 5’pB: 5’ piggyBac end, 3xP3: three tandem repeats of Pax-6 homodimer binding site fused to a basal promoter element, ECFP: enhanced cyan fluorescent protein, attB: bacterial attachment site, TRE: tetracycline response element, AmC: *Anemonia majano* cyan fluorescent protein, attL: *attP*+*attB* hybrid site, attR: *attB*+*attP* hybrid site.

**Table 1.** Injections of attB-TRE-AmCyan into three lines (A, D and C) carrying attP-AeCPA-tTA. The survival rate was calculated as: (adult G_0_ males+adult G_0_ females)/injected eggs, and the minimum transformation efficiency as: positive G_1_ pools/(adult G_0_ males+adult G_0_ females). Chr.: chromosome.

| **Target line** | **Eggs** | **G<sub>0</sub> larvae** | **Adult G<sub>0</sub> males** | **Adult G<sub>0</sub> females** | **Survival** | **Pools(positive G<sub>1</sub>)=sex of G<sub>0</sub>** | **Minimum transformation efficiency** |
| --- | --- | --- | --- | --- | --- | --- | --- |
| A<br>(Chr. 1) | 318 | 25 | 11 | 9 | 6.29% | A(0),B(0)=fem<br>C(1),D(9),E(0)=male | 10% |
| D<br>(Chr. 2) | 334 | 21 | 10 | 8 | 5.39% | A(28),B(14)=fem<br>C(5),D(1)=male | 22% |
| C<br>(Chr. 3) | 329 | 11 | 3 | 8 | 3.34% | A(2),B(0)=fem<br>C(86)=male | 18% |

The target lines carry the transgene attP-AeCPA-tTA (Figure 8b), which uses a 1.6 kb fragment the *Ae. aegypti Carboxypeptidase A* promoter (AeCPA) to express tTA (Gossen and Bujard, 1992). The interaction, if any, between the AeCPA-tTA and TRE-AmC elements (represented as grey segments in Figure 8b-e) is not relevant for this study and is not discussed.

We injected between 318 and 334 eggs and made between three and five G_0_ pools per target line (Table 1). For line A we recovered positive G_1_ (i.e., individuals bearing both transformation markers) in two out of five pools, for line D we recovered positive G_1_ in all four pools, and in line C we recovered positive G1 in two out of three pools. We established the lines by crossing the positive G_1_ from each pool separately with wild-type counterparts, and then collected G_2_ samples for analysis of the insertion site.

After genomic DNA extraction from the different G_2_ lines, we corroborated the insertion of the transgene in the target site by amplifying and sequencing both junctions (Figure 8d, Supplementary Table 13). We used primers 6707 and 6799 (Figure 8d, green arrows) to amplify *attL*, expecting a product of 1101 bp, and primers 6795 and 6791 (Figure 8d, yellow arrows) to amplify *attR*, expecting a product of 1393 bp. All lines produced amplicons of the expected sizes (Figure 8f), and all sequences showed hybrid *attL* and *attR* sites (Supplementary Table 13). The PhiC31 integrase plasmid itself carries an *attB* site, a relic of its construction history. This may have lead to some integration of the PhiC31 plasmid itself. The PhiC31 integrase plasmid has no fluorescence marker so such insertions would not have been recognised; since integration into *attP* is exclusive (first insertion alters the site) double insertions were not expected or identified. To the extent that some wild-type chromosomes were modified in this way and therefore not accessible to integration by the marked plasmid, the insertion rates of Table 1 may be slight underestimates.

## 4. Discussion

Both Cre and PhiC31 catalyse recombination between specific sequences, *lox* and *att* sites, respectively. These sequences are long enough to be absent from most genomes, so that recombination can be specifically directed to artificially integrated sequences. Furthermore, these sequences are stable in the absence of the relevant enzyme, and the enzyme is essentially inert in the absence of the target sequences. This allows considerable scope for controlling the recombination reaction by combining the two components only under specific conditions. Examples might include crossing two strains, injecting a source of enzyme, e.g. a plasmid encoding the enzyme, etc.

Site-specific integration is desirable over other methods as it can use target sites that have previously been characterised for their pattern and levels of expression, avoiding having to maintain and characterise several mosquito lines with different insertions, or even having to resolve multiple insertions. Targeted integration may now be achieved using CRISPR/Cas-mediated homologous recombination in many cases. This method requires, however, knowledge of the target genome to be able to generate suitable guides (followed by analysis of the efficiency/specificity of those guides) and homology arms, and identify desirable target sites avoiding repetitive sequences and pseudogenes. It also depends on homology-directed repair (HDR) to happen preferably over non-homologous end joining (NHEJ) after the double-strand break is made by the Cas enzyme. Although this may be achieved by silencing elements of the NHEJ pathway (see Basu *et al*., 2015 for example in *Ae. aegypti*), this adds an extra layer of complexity to the process.

Considering all the potential limitations described above for CRISPR/Cas-mediated integration, PhiC31 remains an efficient alternative. Unlike for CRISPR/Cas, one *att* site needs to be integrated by other means before the PhiC31 method can be used for integration into the germline. This may be efficiently achieved by establishing “docking lines” where *attP* is inserted in a relatively well-characterised site. This characterisation of the site can be done by sequencing (if a reference genome is available) or by analysing the fitness of the transgenic individual (if the genome is unknown). Besides, although we only inserted a plasmid of approx. 6 kb, it is known that this recombinase can integrate the over 40 kb *Streptomyces* phage PhiC31 genome (Chater et al., 1981), and it is thought that it could have no upper size limit (Olivares and Calos, 2004), broadening the spectrum of possible transformations.

Using a plasmid-encoded source of PhiC31, we achieved efficient integration into *attP* sites in each of the three chromosomes of *Ae. aegypti*, with efficiency levels between 10 and 22%. Previous experiments in *Ae. aegypti* by Nimmo *et al*. (2006) using PhiC31 mRNA, had an average efficiency of 23%. It is worth noting the following differences between our and their study: i) we injected 318-334 eggs while they injected 689-1195; this affects the number of plasmids that can be injected per session, or the number of people/equipment necessary to perform the task, as well as making more difficult the handling of the G_0_ depending of the survival rate, ii) we obtained in total between 10 and 88 positive G_1_ in both male and female G_0_ pools, while they obtained between 1 and 11 positive G_1_ males only in the male G_0_ pools (no information is provided regarding G_1_ females); although theoretically one single transgenic G_1_ individuals is required to start a line, as initial survival of transgenics can be poor, the generation of more than one transgenic G_1_ is desired, iii) we used three mosquito lines with single *attP* sites in each chromosome of *Ae. aegypti*, while they used two lines with single insertions, one with two insertions, and one with four insertions. Even if the efficiencies are similar, it is clear that the use of a plasmid for the expression of the PhiC31 integrase simplifies the microinjection procedure.

PhiC31 integrates the entire plasmid but, as we demonstrated, undesired sequences can be flanked by *lox* sites to allow efficient subsequent removal. In one special case of this, the docking site might incorporate both *attP* and *lox*, allowing cassette exchange in two steps (PhiC31 recombinase and Cre provided sequentially) or one step (PhiC31 recombinase and Cre provided together) (Haghighat-Khah et al., 2015).

Site-specific recombination has many potential uses, and has been used extensively in insects genetics (e.g., Venken *et al*., 2016; Ahmed and Wimmer, 2022). Potentially deleterious transgenes may be inactivated by inserting a redundant fragment of DNA that interrupts the gene, then activated by recombination-induced removal of the redundant DNA post-integration. Recombination sites may be inserted within or between genes, allowing specific chromosome deletions or rearrangements to be generated as desired.

We also demonstrated that more complex rearrangements can be induced post-integration. Using overlapping *lox*P and *lox*N pairs, we were able to recover both *lox*P and *lox*N recombinants. These are mutually exclusive pathways, given that recombination between *lox*P removes one of the *lox*N sites, and *lox*N recombination similarly removes the *lox*P site.

Post-integration Cre-*lox* manipulations are preferably managed by crossing in a source of Cre. This allows the use of very low numbers of insects, such as may be available in the first generations after constructing a new transgenic line, whereas injection experiments need substantial numbers of eggs, as well as being considerably more time-consuming than simple crosses. It also removes the effect of injection as a confounding factor when comparing pre- and post-recombination phenotypes. We demonstrated that Cre-mediated *lox*P recombination can be efficiently achieved in the germline using an integrated Cre source (Shu-Cre). With recombination rates at the observed level – approx. 15-17% with two types of *lox* sites present and 32-37% with *lox*N only, though this may vary somewhat by insertion – the desired recombinant can likely be recovered from the progeny of only one or a few initial Cre-*lox* insects.

This set of experiments expands the toolbox for synthetic biology in *Ae. aegypti*. It is highly likely that these methods can be transferred to other mosquito species, as many species of public health importance are now reared and manipulated in the laboratory. In turn, this expands the options both for fundamental studies to better understand the insects, and for applied approaches aiming to develop new methods of controlling mosquito populations and the pathogens that they transmit.

## 6 Author Contributions

LZCP, PYLT and LA designed the research. LZCP, RW, PYLT, VD, EK, PC, WL, SR, MN, AU and LW performed the research and maintained the mosquito colonies. LZCP, PYLT, SB and PL designed the plasmids. MAEA contributed reagents. LZCP prepared the first draft and all authors contributed to reviewing and editing of the manuscript.

## 8 Acknowledgments

The authors thank Victoria Sy for her assistance in the Insectary during the development of the attP-AeCPA-tTA transgenic line.

## 10 Data Availability Statement

All the data collected is presented in Supplementary Material.

