## Supplemental information for "Optimizing CRE and PhiC31 mediated recombination in *Aedes aegypti*"

### 1 Supplementary Figures and Tables

#### 1.1 Supplementary Figures

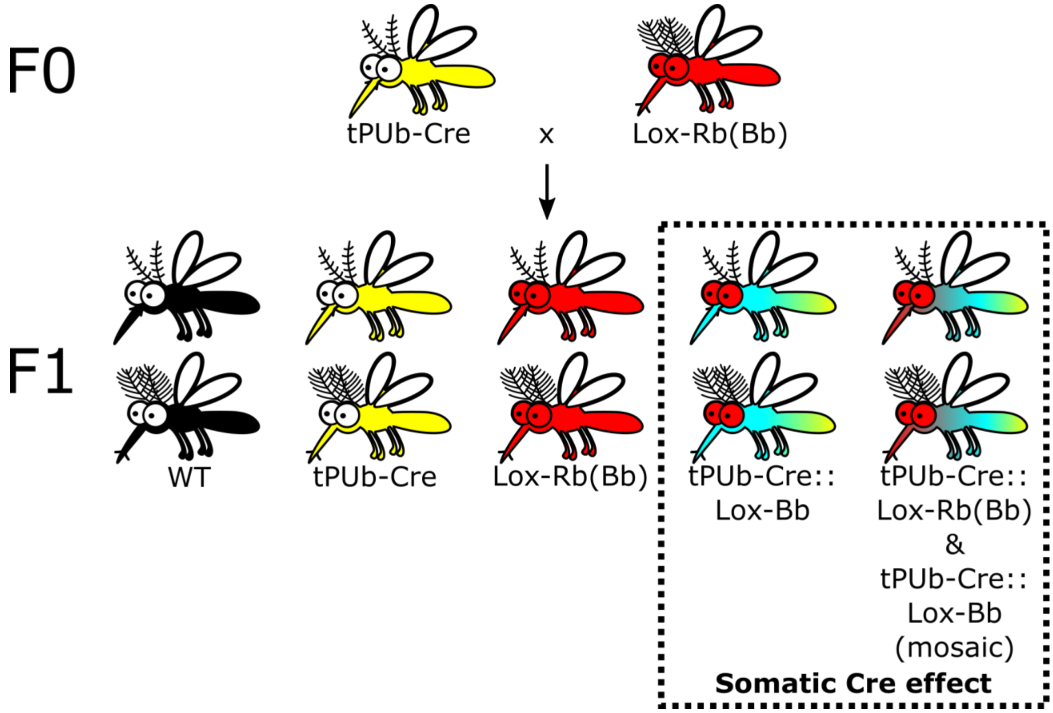


**Supplementary Figure 1. Schematic representation of the expected phenotypes of a cross between a hemizygous line expressing Cre recombinase ubiquitously (tPUb-Cre) and a hemizygous line carrying construct PUb-loxP-mCh-stop-loxP-AmC (Lox-Rb(Bb)).** The expected phenotypes of the $F_1$ include: wild-type (WT), single hemizygous carrying tPUb-Cre, single hemizygous carrying PUb-loxP-mCh-stop-loxP-AmC (Lox-Rb(Bb)), double hemizygous carrying tPUb-Cre and the recombined version of PUb-loxP-mCh-stop-loxP-AmC without the whole body red marker and expressing blue instead (tPUb-Cre::Lox-Bb), double hemizygous as before but carrying non-recombined versions as well (mosaics).

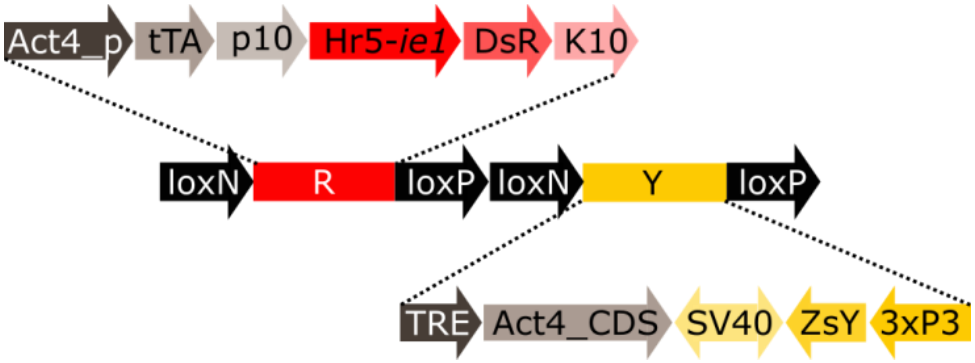


**Supplementary Figure 2: Schematic representation of the complete plasmid loxN-R-loxP-loxN-Y-loxP.** Act4_p: *Aedes aegypti Actin-4* promoter, tTA: tetracycline-controlled transactivator, p10: *Autographa californica* nucleopolyhedrovirus (AcMNPV) *p10* 3'UTR, Hr5-*ie1*: AcMNPV *ie1* promoter fused with homologous region 5 (hr5) enhancer, Shu_p: *shut-down* promoter, DsR: *Discosoma* sp. red fluorescent protein, K10: *Drosophila melanogaster* K10 3'UTR, TRE: tetracycline response element, Act4_CDS: *Ae. aegypti Actin-4* coding sequence, SV40: Simian virus 40 PolyA sequence, ZsY: *Zoanthus* sp. yellow fluorescent protein, 3xP3: three tandem repeats of Pax-6 homodimer binding site fused to a basal promoter element.

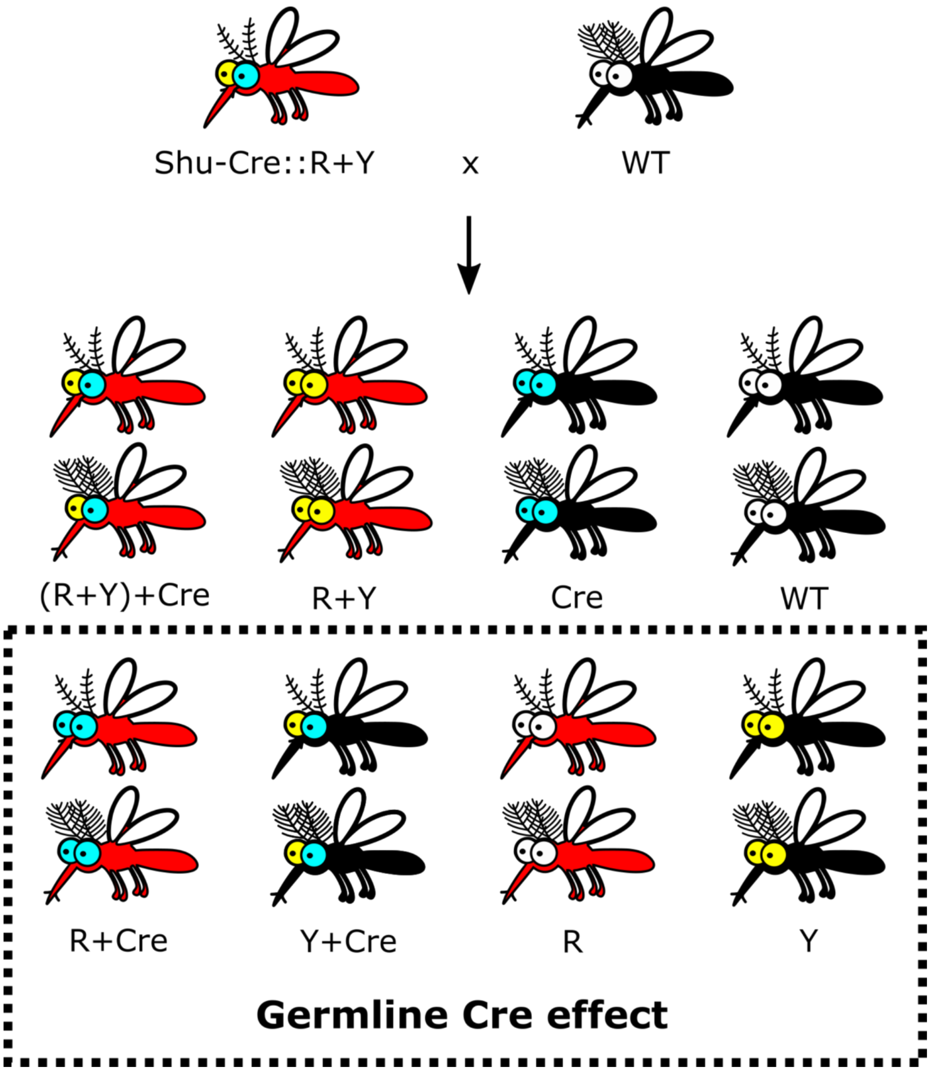


**Supplementary Figure 3:Schematic representation of the expected phenotypes of a cross between double hemizygous expressing Cre recombinase in the germline (Shu-Cre) and carrying the reporter construct loxN-R-loxP-loxN-Y-loxP (R+Y), and wild-type individuals (WT).** The expected phenotypes include: double hemizygous carrying loxN-R-loxP-loxN-Y-loxP and Shu-Cre ((R+Y)+Cre), single hemizygous loxN-R-loxP-loxN-Y-loxP (R+Y), single hemizygous Shu-Cre (Cre) and wild-type (WT), as well as the recombined genotypes carrying loxN-R-loxP and Shu-Cre (R+Cre, removal of segment Y), only loxN-Y-loxP and Shu-Cre (Y+Cre, removal of segment R), only loxN-R-loxP (R), and only loxN-Y-P (Y).

### 1.2 Supplementary Tables

**Supplementary Table 1:** Genbank accession numbers of the plasmids/transgenes.

| Plasmid/transgene name | Accession number |
|---|---|
| AGG1561_tPUb-Cre | OR242269 |
| AGG1959_Shu-Cre | OR242275 |
| AGG1572_PUb-loxP-mCh-stop-loxP-AmC | OR242270 |
| AGG1600_loxN-R-loxP-loxN-Y-loxP | OR242271 |
| AGG1741_A-loxN-B-loxN-C | OR242273 |
| AGG1029_attP-AeCPA-tTA | OR242268 |
| AGG1733_AePUb-PhiC31 | OR242272 |
| AGG1755_attB-TRE-AmC | OR242274 |

**Supplementary Table 2:** Insertion sites of the lines used in the experiments.

Sequences of the insertion sites of the lines used in the experiments, obtained by flanking PCR.

| **Transgenic line** | **5' insertion site** | **3' insertion site** | **Genome location (Chromosome: from-to)** |
|---|---|---|---|
| tPUb-Cre | GATCCATTATTACAATTGCCAAAATTATTACAATTGTGCAATTTGAAGTAGATTATAAAACTGAATTTAGTATAATACCATTAAATTCCACTAGAGTTTGTATCCTTTGACAGATACGCGTATTTCGACCTCAACTGTAAGGCCGTCTTCAGTGTCGTGTAATAGACTCGACTGTGCAATTTTGTTACATGTGATAATTTTCGTCATCGAAGTTTTCCCATAAAATATGGGAAATCAAAGACGCTGTTTGAAATGCCACTCTATTTTACAATCGTTGTGAAATGTTGAGCACTCATCCCAATGGCACTGCAGCAGCGTCCTCGAAAGCATCCTCAAATTGCCGTATTGTCAATTATTCGTAATAAATCGGTTATTGAATGATGCTACGAGATGAATAAAGAGATGTGGTAAACACATTAGGGAATATTACACAATCATCTCTT**TTAA** | **TTAA**TCTAGCTGGTGGCACTTTTCGGGGAAATGTGCGCGGAACCCCTATTTGTTTATTTTTCTAAATACATTCAAATATGTATCCGCTCATGAGACAATAACCCTGATAAATGCTTCAATAATATTGAAAAAGGAAGAGTATGAGTATTCAACATTTCCGTGTCGCCCTTATTCCCTTTTTTGCGGCATTTTGCCTTCCTGTTTTTGCTCACCCAGAAACGCTGGTGAAAGTAAAAGATGCTGAAGATCAGTTGGGTGCACGAGTGGGTTACATCGAACTGGATCTCAACAGCGGTAAGATCCTTGAGAGTTTTCGCCCCGAAGAACGTTTTCCAATGATGAGCACTTTTAAAGTTCTGCTATGTGGCGCGGTATTATCCCGTATTGACGCCG | Unknown |
| Shu-Cre | GATCCTTTCTCTTTTTGGCGATTTTTCAAAAGCAGAAAAGATGCGCAGATAATCCCATCCGAAATGTCGTATGGCGACTCA**TTAA** | **TTAA**GATACAACATCGAAGGGTCTTTCGAACTTTTCACCGACTACAGCGGGGATTCGCTAGTTGGAACACGACTGCCTTCCATCTAACGAATCCGACCCGTTAGGTGGAACGACTGACAGCTGGTGAAAATGCTCCAGACTAACGCATCCGGAACGCAAATCGTCACGAAACGCACATATACATACACATTTGTTCTATATATGGAGATTTCAATGCGACGGTACACAGCCAATAG | 1: 148,452,176-148,452,574 |

| | | | |
|---|---|---|---|
| | | TGGTTTCTGCGCTCGTGTACATACACG TTACATGTCACCAACACTACTTCAAGA ATGTTGGTTGAAAATATAAACAAAGA TC | |
| PUb-loxP-mCh-stop-loxP-AmC | GATCTTCCAGAAACGATTTGAGGA AGTATGGGAGTTTACGTATTCTCGG CAATAAAGTCCCTAGTAAAATTATC AAATGTCAAAACTAAAAATGCAGT TTTCTCCGTAATAATTACCAAATCT GGGAAAATAAAAACACACATTCTT TCCACAGACAAACAGACGTAACAG TTAGAACAAATTTATTCATATTCCA TCGCCCAGTTTACACCATCATCTGG TGAGCATGTTACACGAAAAGGTGT TTCGTGTAACGAGACCGGCAGATG GCGGTAGTGTGAAACGTCAAACGC GAAGAAAAACGATGCGCGCGCCTC TGGGTGTGAGATTTGTAACATAGCG AATTTGAAATGATTG**TTAA** | No useful bands obtained | 3: 337,156,953-337,157,313 |

| | | | |
|---|---|---|---|
| loxN-R-loxP-loxN-Y-loxP | GCAACCCCAAATCTACTAGAGACA<br>CATTATTTGATCCTTGAGACACGCA<br>CAAATGTCGTTTTTGTTGCGCTGTG<br>TAATGGTAACCCCTAATTTTCAGAT<br>AAATCAGAACAGGTCTCCTTCTTCT<br>TTCCTTGGCTCCCTTCAACATACAT<br>TACCTATTTCCTACCACTTTGTAGC<br>TAGTTCCTATAAACATGTATGCAAG<br>TTCTGTCATCTTGCTCGCAAGGTAA<br>TATAGATTGATTGACAGATAGGCA<br>ACGGCTCTTATGTGTTTTACTAATT<br>TATATTTGTTTATCACATGATATGA<br>TGTACAAAACCATACTAATCGAAG<br>CTTTTTTCTTTTCTCCTTTTCAGGTA<br>TGGACGAAATTAGCAAAACATAAT<br>TAAGGTACGGTTAAAAATGTCTGA<br>AAACAAAATATATTACGGATTAAA<br>TATTAAGGAGGCCCTTTATCA**TTAA** | **TTAA**ATTATTATCATGCTTTAAAGTTT<br>CGCCTCTTGCTCGTGCTAACTGCTACT<br>GGAACAGTTCAGAGTTTGAACCATGC<br>AACAAGTTTTTCTCTAAACATTATTCG<br>AAACTAGTTATGGAAACGAATATTTA<br>TAGAAAGCATTTCGATATAAACGTGT<br>ATGTCTCATATCGCAATGGAAAGCTTC<br>GACTTACATATAAATTATAACAATTA<br>ACACGTTACTTTCCTAGTTTTCTTTTTA<br>GTTGGTTTGACTATCTCCAACCACAAT<br>GATTTCCGCTGATTTTTCTATTCCAAT<br>AGTTTTCGCTTCAACGATACTCAGCAT<br>TCGAGCGATTCTAACCATTATACAATC<br>ACAATTTCTGTGACACAAGCACCCGT<br>CCCATCTAGTGCTATAAGTATTTTCTA<br>GTAATATTCCATGTGACTTGACTACTG<br>AACGTGTTGTGCACACTTACAAACTGT<br>AAACTATATATACTTACTTAAACAGCC<br>TAGCGTCAAACAGGCTACCCCGTCTG<br>CCATATCTGCGCCATCCAGCGATC | 2: 316,878,424-316,879,695 |
| A-loxN-B-loxN-C Line D | AGTCGCTCCGATCCGATCCGTTGTT<br>GATTTGGCCGAAAGAATTCCGCTG<br>AATCTTTCATCTCCATCCATGTTCG<br>GAGTTCAGAGTTGGAGAAAAATCA<br>AAGTACGTGATAAAATTCTAAAAA<br>CAACACAATTCCAATGTTTATTCCA<br>ACGTGGACTTTGAGCAGGAAATAG<br>ATTCCGTAAAAGTTTTGTAATAATA<br>GATCTTGCCACTTTCT**TTAA** | **TTAA**GATTCATCCGCCCGCGTGTCGTC<br>GTCGTTTGACTTACACACAGCGTCTTC<br>CCCTTGCGTATCTCCCGTATCTTCTCG<br>ACGATTTGTCGGTTAGAAAAATGACC<br>TCCGTGATGCTCTTTCTCCCGGAGTCG<br>GAGCTGGNATTTAAACACATTCTGTA<br>AACAGATAAGAAATGGGAAT | 2: 255,619,007-255,619,397 |

| | | | |
|---|---|---|---|
| A-loxN-B-loxN-C<br>Line E | No useful bands obtained | **TTAA**AATTAATTTTATCTATTATCTTA<br>CTTGAACATAGAGATTAATCTGCTCG<br>GTGATCAACACCGAGACGCTTTTGTA<br>CTTTTTGTTGATCTTGGTACCGAGCGT<br>CATAAAACGCAACAGGAACGTTCCAA<br>ATGCACAGGACCACACCAGTGCCTCC<br>AAGTTGAATTGCGAGTGGAGATGAAC<br>AGAGTCCTGCAACGGGAAATAATAAA<br>TACGAGCCTAATTAGCATTCCAAATA<br>CAAACCTTCAACCACTCGACACTGAG<br>GAATGCCAACAATAGCAGTGTTATGA<br>TAAATGCAGCCGAAACGATCACGTCC<br>ACCGAGTTCTGTGGTCCACGTCGTTTC<br>AGATAGGAGCGCACACACAGCCAGGT<br>CTTGATGTCCGCAC | 2: 127,489,035-<br>127,489,416 |
| attP-AeCPA-tTA<br>Line A | CGGGTTGGCTGTTGATATGTTGGAG<br>AAACTGTCCACACAGGCTTGTATAT<br>TGAACAGATGATAAATAGTTTAATT<br>TATTTTTCAAGCCTTCTGCTGATCTT<br>GTATTTTTGTTTTGCTAAATCGTGG<br>GTAAGACAATCCTTTGACTGTCCTA<br>CCCAGGTATTTTTGTGCGTAGTACA<br>TGTCCTACCTGTCCTACCCACTTCC<br>CGCGCCACTGGTTCCCTTTATTCTTT<br>GGAACAATGAACAATATTTAATTTG<br>TGTCTGAAATTTACCCAATTTAATA<br>TTCGTACACCATTTTACACTTACAA<br>AATTTAAAAATGCAATCATATTATT<br>ATTTTTCTAAACATTTATATCGTTGT<br>TTCGATTTATTAAAGTACATTTCTC<br>GCAAGATTTATGATCTCGCCCGAAA<br>AGCCTTCGGTAATCAAGTCTACAGC<br>TTGTTGTTATATTATGTTGTTATAGT<br>ATACTGAGAAGATCCTCCTCTGTCT | **TTTAA**GTCGCTGTAAATGCTTCTTCAT<br>CCAAAAGCATGATCGATACTTACAAA<br>ACACTACTGCAAGCGTGTGTGAGAGA<br>GAGTCTGGCACGAATGCTGCTTTACC<br>GCGGCGACGAGTAAAGCAGATTGTCC<br>AGGTTAAGCTTGTTCTTGTTGGGAAGC<br>GGTGCTCCGTGCCCTGGCCGGCCCCA<br>GAAGCTGGACCACGTGTCGAAGTGCT<br>GTTTGCTGGTTTCGAACTCCACCACTC<br>GGGTGGTTTCCATTCGGCGCTGCTTCT<br>GGTTCATTTGGGTGCAAAGCTCGTGC<br>AGGTATTCATCTCCCGAGTGCCGTGCC<br>GCTACGCTGAAAGCAGGTGGAAGAAA<br>GCGGTGTGGGATGGTTAGTTCCGAAT<br>TGGCCTGTTGGGGACTACTGAATATTA<br>GGGCTGTACTATTTGCTGAATTTACTA<br>ATTAGTGGTGCCGTCGCTCTAGGGGG<br>ATGATAGTTTACCCATTGGAGCTGTAC<br>TTATCGATTTTGGTAACGTCGGTCGTC | 1: 25,788,700-25,789,979 |

| | | | |
|---|---|---|---|
| | CACAGTAGTGATAGTTTTTAATAAA<br>TGGCAAATTAATGCGTATCGAATG<br>ATAGATGGCATCCACGACTTTCATT<br>TTTGATAAAATTTTTCAGTTTGCGT<br>ACATCATTGATGGTTTTTCACCAAC<br>TTGATGTGGAAGCAGTGCCCCGTAC<br>GAATATGAA**TTTAA** | CCGAGGCTAATGGGTCTCGGTTGGGA<br>GCGGCGTTTGGCCAGCAGCGGGACCA<br>ATTCCACACCTCCG | |
| attP-AeCPA-tTA<br>Line C | CTGTATAATATAGTAGAAAATACA<br>ATATGCTTCTTTTACTCAGTTCTCAC<br>TTGCATGCCTTAACGC**TTAA** | **TTAA**ACCTAAACTGCAACGAAAGGTT<br>ATCTCCTCAATATGATTTTCCTCTTCCT<br>TAAAATACATCTCATTCGATTCCGCTC<br>ACTCATTCCTAATATGGCTGCTTGCAC<br>TTTACAATTAATGTATGCATGTATGTA<br>TTTATGTGTGTTGCATATCGTTACAAA<br>CTAGTTAGTTCACTTTGCTGACTACAA<br>ACTACAAAAACTGAGACTTGATAATC<br>ATCTGGCTGAAAATCTATACATTTCGA<br>AAGAATAAC | 3: 375,528,567-<br>375,528,883 |
| attP-AeCPA-tTA<br>Line D | CATGGATTCAATGGTGGACGAAGG<br>CGGTCGATGGGCGGTGACACAATC<br>CGGACGAGACCCGATGGCAATAGC<br>GGGCCACCGGCGGTGTACGTTGAT<br>GGCGGCGAAGAACAAGAGTCCCGA<br>CGGGGAACTGGTCGCTTGATGGAA<br>TGGCGACGGAGGCAATGGCGGTGG<br>ACTACGGAGGCACTGGGGCTTGGA<br>AGATGGCGGTGGTGTAACGGAACG<br>GTCGTTCGTTGACGGTAACGACCCG<br>AATTCTCGGACAGCGATGGCGTCTG<br>GGCTGTCTCTGCCTGGTTGCGCTTC<br>GATTCTGGCCACTTCCGTTCACTCC<br>GTGTCACTAGCACTTTCCAGGAACT<br>CGTCTGCCGAAGAAAAAACTCAAA<br>AGGAGAAGAAAAAGGCCCGCTTCG<br>TGGACCTGCACACAAAATTAGTAC<br>CCTGTGACGTCGCATATTTTAGCCT | **TTAA**GCAGCTTTGACTATTGACTAATA<br>GACTAACAATACTAACAATTGTCTATT<br>GTCCAACTAACAATCACCACCTATCTA<br>CAAAGAAACAAAAGCACTAATTATTA<br>ACACTTCCACTTTTCTTATTACTTTACT<br>TCTAACCTTTGTAAAAAACTTTCAACT<br>AACAACGCGCACTTCTTATCACTTTCA<br>GATTTTCCAGGCAAAAGAGATCTTTCT<br>TAGCAGTTTCAAACTATTTATAAGAAC<br>TTAAAATTTATCAGTGGAGCCGAACT<br>ATCTAAAATCGTTCGTGCTTAAAAGTC<br>ACGAACGATTTTTGACAGTTATGCTCA<br>TAACATGGCGGTTTACCCCAATAGCG<br>ATTTGTGCTCCCCCTATGAGCGAATAG<br>TACATATCCG | The insertion site matches three locations within chromosome 2 with 99% identity:<br>2: 14,711,505-14,712,517<br>2: 138,885,258-<br>138,886,271<br>2: 108,726,025-<br>108,727,036 |

| | |
|---|---|
| | CTAACAAACTAGCAACTATTCACA<br>AAACATTAAATTACCTTTTTCTTGT<br>GTTTGTCTTAATATAACTGTTGTTTC<br>ACACTGTTTCGTATGTTTTTAGCGT<br>GTTATCTATAATTGGTTTTATTCAC<br>ATTAGCTTTAGGAAACAATGATCAC<br>TTTTTGTTACTAACCATACTAATAC<br>TGCTAATACACTAT**<u>TTAA</u>** |

**Supplementary Table 3:** Number of individuals of each phenotype observed in the $F_2$ of crosses between double hemizygous obtained by crossing females hemizygous for the reporter loxN-R-loxP-loxN-loxP and males hemizygous for tPUb-Cre, and wild-type counterparts.

Rep1 to 3 indicate each of the replicates (3x 300 eggs each). The observed values (Obs) were calculated as the average of the values of the three replicates. For statistical analysis the observed values were obtained adding up individuals carrying non-recombined and recombined versions of the reporter construct. The expected values (Exp) were calculated as 25% of the average of the total number of individuals screened in the replicates, considering complete segregation of constructs and full survival of the double hemizygous, each of the single hemizygous, and wild-type (WT). Phenotypes in bold represent the recombined versions.

| | **loxN-R-loxP-loxN-Y-loxP::tPUb-Cre** | | | | | | | | | |
|---|---|---|---|---|---|---|---|---|---|---|
| | **x WT female** | | | | | **x WT male** | | | | |
| | **Rep1** | **Rep2** | **Rep3** | **Obs** | **Exp** | **Rep1** | **Rep2** | **Rep3** | **Obs** | **Exp** |
| loxN-R-loxP-loxN-Y-loxP::tPUb-Cre | 48 | 46 | 51 | 48 | 55 | 53 | 61 | 70 | 61 | 64 |
| **loxN-R-loxP::tPUb-Cre** | 0 | 0 | 0 | 0 | | 0 | 0 | 0 | 0 | |
| **loxN-Y-loxP::tPUb-Cre** | 0 | 0 | 0 | 0 | | 0 | 0 | 0 | 0 | |
| loxN-R-loxP-loxN-Y-loxP | 64 | 56 | 46 | 55 | 55 | 72 | 61 | 74 | 69 | 65 |
| **loxN-R-loxP** | 0 | 0 | 0 | 0 | | 0 | 0 | 0 | 0 | |
| **loxN-Y-loxP** | 0 | 0 | 0 | 0 | | 0 | 0 | 0 | 0 | |
| tPUb-Cre | 58 | 60 | 49 | 56 | 55 | 54 | 61 | 68 | 61 | 65 |
| WT | 52 | 64 | 67 | 61 | 55 | 56 | 63 | 85 | 68 | 65 |
| Total | 222 | 226 | 213 | 220 | 220 | 235 | 246 | 297 | 259 | 259 |
| | $X^2$= 0.38606, df= 3, p= 0.9431 | | | | | $X^2$= 0.79508, df = 3, p= 0.8506 | | | | |

**Supplementary Table 4:** Number of individuals of each phenotype observed in the $F_2$ of crosses between double hemizygous obtained by crossing females hemizygous for tPUb-Cre and males hemizygous for the reporter loxN-R-loxP-loxN-loxP, and wild-type counterparts.

Rep1 to 3 indicate each of the replicates (3x 300 eggs each). The observed values (Obs) were calculated as the average of the values of the three replicates. For statistical analysis the observed values were obtained adding up individuals carrying non-recombined and recombined versions of the reporter construct. The expected values (Exp) were calculated as 25% of the average of the total number of individuals screened in the replicates, considering complete segregation of constructs and full survival of the double hemizygous, each of the single hemizygous, and wild-type (WT). Phenotypes in bold represent the recombined versions.

| | **tPUb-Cre::loxN-R-loxP-loxN-Y-loxP** | | | | | | | | | |
|---|---|---|---|---|---|---|---|---|---|---|
| | **x WT female** | | | | | **x WT male** | | | | |
| | **Rep1** | **Rep2** | **Rep3** | **Obs** | **Exp** | **Rep1** | **Rep2** | **Rep3** | **Obs** | **Exp** |
| tPUb-Cre::loxN-R-loxP-loxN-Y-loxP | 49 | 57 | 50 | 52 | 60 | 60 | 71 | 68 | 66 | 66 |
| **tPUb-Cre::loxN-R-loxP** | 0 | 0 | 0 | 0 | | 0 | 0 | 0 | 0 | |
| **tPUb-Cre::loxN-Y-loxP** | 0 | 0 | 0 | 0 | | 0 | 0 | 0 | 0 | |
| loxN-R-loxP-loxN-Y-loxP | 55 | 55 | 63 | 58 | 59 | 74 | 70 | 57 | 67 | 66 |
| **loxN-R-loxP** | 1 | 0 | 0 | 0* | | 0 | 0 | 0 | 0 | |
| **loxN-Y-loxP** | 0 | 0 | 0 | 0 | | 0 | 0 | 0 | 0 | |
| tPUb-Cre | 37 | 61 | 62 | 53 | 59 | 68 | 55 | 56 | 60 | 66 |
| WT | 66 | 72 | 84 | 74 | 59 | 64 | 63 | 85 | 71 | 66 |
| Total | 208 | 245 | 259 | 237 | 237 | 266 | 259 | 266 | 264 | 264 |
| | $X^2$= 0.47571, df= 3, p= 0.9242 | | | | | $X^2$= 2.5931, df = 3, p= 0.4587 | | | | |

*Rounded down from 0.33.

**Supplementary Table 5:** Percentage of individuals of each phenotype observed in the $F_2$ of crosses between double hemizygous obtained by crossing females hemizygous for the reporter loxN-R-loxP-loxN-loxP and males hemizygous for tPUb-Cre, and wild-type counterparts.

Rep1 to 3 indicate each of the replicates (3x 300 eggs each). WT: wild-type. Ave: average percentage amongst the three replicates. S.E.: Standard error. Phenotypes in bold represent the recombined versions.

| | **loxN-R-loxP-loxN-Y-loxP::tPUb-Cre** | | | | | | | | | |
|---|---|---|---|---|---|---|---|---|---|---|
| | **x WT female** | | | | | **x WT male** | | | | |
| | **Rep1** | **Rep2** | **Rep3** | **Ave** | **S.E.** | **Rep1** | **Rep2** | **Rep3** | **Ave** | **S.E.** |
| loxN-R-loxP-loxN-Y-loxP::tPUb-Cre | 21.62 | 20.35 | 23.94 | 21.97 | 1.05 | 22.55 | 24.80 | 23.57 | 23.64 | 0.65 |
| **loxN-R-loxP::tPUb-Cre** | 0.00 | 0.00 | 0.00 | **0.00** | 0.00 | 0.00 | 0.00 | 0.00 | **0.00** | 0.00 |
| **loxN-Y-loxP::tPUb-Cre** | 0.00 | 0.00 | 0.00 | **0.00** | 0.00 | 0.00 | 0.00 | 0.00 | **0.00** | 0.00 |
| loxN-R-loxP-loxN-Y-loxP | 28.83 | 24.78 | 21.60 | 25.07 | 2.09 | 30.64 | 24.80 | 24.92 | 26.78 | 1.93 |
| **loxN-R-loxP** | 0.00 | 0.00 | 0.00 | **0.00** | 0.00 | 0.00 | 0.00 | 0.00 | **0.00** | 0.00 |
| **loxN-Y-loxP** | 0.00 | 0.00 | 0.00 | **0.00** | 0.00 | 0.00 | 0.00 | 0.00 | **0.00** | 0.00 |
| tPUb-Cre | 26.13 | 26.55 | 23.00 | 25.23 | 1.12 | 22.98 | 24.80 | 22.90 | 23.56 | 0.62 |
| WT | 23.42 | 28.32 | 31.46 | 27.73 | 2.34 | 23.83 | 25.61 | 28.62 | 26.02 | 1.40 |

**Supplementary Table 6:** Percentage of individuals of each phenotype observed in the $F_2$ of crosses between double hemizygous obtained by crossing females hemizygous for tPUb-Cre and males hemizygous for the reporter loxN-R-loxP-loxN-loxP, and wild-type counterparts.

Rep1 to 3 indicate each of the replicates (3x 300 eggs each). WT: wild-type. Ave: average percentage amongst the three replicates. S.E.: Standard error. Phenotypes in bold represent the recombined versions.

| | **tPUb-Cre::loxN-R-loxP-loxN-Y-loxP** | | | | | | | | | |
|---|---|---|---|---|---|---|---|---|---|---|
| | **x WT female** | | | | | **x WT male** | | | | |
| | **Rep1** | **Rep2** | **Rep3** | **Ave** | **S.E.** | **Rep1** | **Rep2** | **Rep3** | **Ave** | **S.E.** |
| tPUb-Cre::loxN-R-loxP-loxN-Y-loxP | 23.56 | 23.27 | 19.31 | 22.04 | 1.37 | 22.56 | 27.41 | 25.56 | 25.18 | 1.42 |
| **tPUb-Cre::loxN-R-loxP** | 0.00 | 0.00 | 0.00 | **0.00** | 0.00 | 0.00 | 0.00 | 0.00 | **0.00** | 0.00 |
| **tPUb-Cre::loxN-Y-loxP** | 0.00 | 0.00 | 0.00 | **0.00** | 0.00 | 0.00 | 0.00 | 0.00 | **0.00** | 0.00 |
| loxN-R-loxP-loxN-Y-loxP | 26.44 | 22.45 | 24.32 | 24.41 | 1.15 | 27.82 | 27.03 | 21.43 | 25.43 | 2.01 |
| **loxN-R-loxP** | 0.48 | 0.00 | 0.00 | **0.16** | 0.16 | 0.00 | 0.00 | 0.00 | **0.00** | 0.00 |
| **loxN-Y-loxP** | 0.00 | 0.00 | 0.00 | **0.00** | 0.00 | 0.00 | 0.00 | 0.00 | **0.00** | 0.00 |
| tPUb-Cre | 17.79 | 24.90 | 23.94 | 22.21 | 2.23 | 25.56 | 21.24 | 21.05 | 22.62 | 1.47 |
| WT | 31.73 | 29.39 | 32.43 | 31.18 | 0.92 | 24.06 | 24.32 | 31.95 | 26.78 | 2.59 |

**Supplementary Table 7:** Number of individuals of each phenotype observed in the $F_2$ of crosses between double hemizygous obtained by crossing females hemizygous for Shu-Cre and males hemizygous for the reporter loxN-R-loxP-loxN-loxP, and wild-type counterparts.

Rep1 to 3 indicate each of the replicates (3x 300 eggs each). The observed values (Obs) were calculated as the average of the values of the three replicates. For statistical analysis the observed values were obtained adding up individuals carrying non-recombined and recombined versions of the reporter construct. The expected values (Exp) were calculated as the 25% of the average of the total number of individuals screened in the replicates, considering full survival of the double hemizygous, each of the single hemizygous and wild-type. WT: wild-type. Phenotypes in bold represent the recombined versions.

| | **Shu-Cre::loxN-R-loxP-loxN-Y-loxP** | | | | | | | | | |
|---|---|---|---|---|---|---|---|---|---|---|
| | **x WT female** | | | | | **x WT male** | | | | |
| | **Rep1** | **Rep2** | **Rep3** | **Obs** | **Exp** | **Rep1** | **Rep2** | **Rep3** | **Obs** | **Exp** |
| loxN-R-loxP-loxN-Y-loxP::Shu-Cre | 41 | 41 | 41 | 41 | 62 | 32 | 43 | 34 | 36 | 66 |
| **loxN-R-loxP::Shu-Cre** | 11 | 7 | 19 | 12 | | 16 | 16 | 13 | 15 | |
| **loxN-Y-loxP::Shu-Cre** | 7 | 6 | 4 | 6 | | 8 | 9 | 2 | 6 | |
| loxN-R-loxP-loxN-Y-loxP | 36 | 38 | 32 | 35 | 61 | 42 | 49 | 36 | 42 | 66 |
| **loxN-R-loxP** | 12 | 15 | 13 | 13 | | 13 | 15 | 26 | 18 | |
| **loxN-Y-loxP** | 9 | 6 | 4 | 6 | | 8 | 6 | 8 | 7 | |
| Shu-Cre | 59 | 62 | 69 | 63 | 61 | 57 | 50 | 64 | 57 | 65 |
| WT | 71 | 70 | 62 | 68 | 61 | 90 | 76 | 74 | 80 | 65 |
| Total | 246 | 245 | 244 | 245 | 245 | 266 | 264 | 257 | 262 | 262 |
| | $X^2$= 0.85374, df= 3, p= 0.8366 | | | | | $X^2$= 2.6204, df= 3, p= 0.4539 | | | | |

**Supplementary Table 8:** Percentage of individuals of each phenotype observed in the $F_2$ of crosses between double hemizygous of the reporter loxN-R-loxP-loxN-loxP and Shu-Cre, and wild-type counterparts.

Rep1 to 3 indicate each of the replicates (3x 300 eggs each). WT: wild-type. Ave: average percentage amongst the three replicates. S.E.: Standard error. Phenotypes in bold represent the recombined versions.

| | **Shu-Cre::loxN-R-loxP-loxN-Y-loxP** | | | | | | | | | |
|---|---|---|---|---|---|---|---|---|---|---|
| | **x WT female** | | | | | **x WT male** | | | | |
| | **Rep1** | **Rep2** | **Rep3** | **Ave** | **S.E.** | **Rep1** | **Rep2** | **Rep3** | **Ave** | **S.E.** |
| loxN-R-loxP-loxN-Y-loxP+Shu-Cre | 16.67 | 16.73 | 16.80 | 16.73 | 0.04 | 12.03 | 16.29 | 13.23 | 13.85 | 1.27 |
| **loxN-R-loxP+Shu-Cre** | 4.47 | 2.86 | 7.79 | **5.04** | 1.45 | 6.02 | 6.06 | 5.06 | **5.71** | 0.33 |
| **loxN-Y-loxP+Shu-Cre** | 2.85 | 2.45 | 1.64 | **2.31** | 0.36 | 3.01 | 3.41 | 0.78 | **2.40** | 0.82 |
| loxN-R-loxP-loxN-Y-loxP | 14.63 | 15.51 | 13.11 | 14.42 | 0.70 | 15.79 | 18.56 | 14.01 | 16.12 | 1.32 |
| **loxN-R-loxP** | 4.88 | 6.12 | 5.33 | **5.44** | 0.36 | 4.89 | 5.68 | 10.12 | **6.90** | 1.63 |
| **loxN-Y-loxP** | 3.66 | 2.45 | 1.64 | **2.58** | 0.59 | 3.01 | 2.27 | 3.11 | **2.80** | 0.26 |
| Shu-Cre | 23.98 | 25.31 | 28.28 | 25.86 | 1.27 | 21.43 | 18.94 | 24.90 | 21.76 | 1.73 |
| WT | 28.86 | 28.57 | 25.41 | 27.61 | 1.10 | 33.83 | 28.79 | 28.79 | 30.47 | 1.68 |

**Supplementary Table 9:** Summary of an ordinary least squares model with the number of individuals observed as the dependent variable and an interaction term between cross and phenotype.

CI: confidence interval. WT: wild-type. Phenotypes in bold represent the recombined versions.

| **Characteristic** | **Beta** | **95% CI** | **p-value** |
|---|---|---|---|
| **Cross** | | | |
| M (WT female x Shu-Cre::loxN-R-loxP-loxN-Y-loxP male) | — | — | |
| F (Shu-Cre::loxN-R-loxP-loxN-Y-loxP female x WT male) | 0.41 | -1.2, 0.40 | 0.3 |
| **Phenotype** | | | |
| Shu-Cre | — | — | |
| **loxN-R-loxP** | -4.3 | -5.1, -3.5 | <0.001 |
| **loxN-R-loxP+Shu-Cre** | -4.5 | -5.3, -3.7 | <0.001 |
| loxN-R-loxP-loxN-Y-loxP | -2.0 | -2.8, -1.2 | <0.001 |
| WT | 0.27 | -0.55, 1.1 | 0.5 |
| loxN-R-loxP-loxN-Y-loxP+Shu-Cre | -1.6 | -2.4, -0.74 | <0.001 |
| **loxN-Y-loxP** | -5.5 | -6.3, -4.7 | <0.001 |
| **loxN-Y-loxP+Shu-Cre** | -5.6 | -6.4, -4.8 | <0.001 |
| **Cross * variables** | | | |
| F * **loxN-R-loxP** | 0.96 | -0.19, 2.1 | 0.10 |
| F * **loxN-R-loxP+Shu-Cre** | 0.84 | -0.31, 2.0 | 0.15 |
| F * loxN-R-loxP-loxN-Y-loxP | 0.97 | -0.18, 2.1 | 0.10 |
| F * WT | 1.1 | -0.02, 2.3 | 0.055 |
| F * loxN-R-loxP-loxN-Y-loxP+Shu-Cre | 0.03 | -1.1, 1.2 | >0.9 |
| F * **loxN-Y-loxP** | 0.63 | -0.52, 1.8 | 0.3 |
| F * **loxN-Y-loxP+Shu-Cre** | 0.46 | -0.69, 1.6 | 0.4 |

**Supplementary Table 10:** Number of individuals of each phenotype observed in the $F_2$ of crosses between double hemizygous of the target lines C or D of A-loxN-B-loxN-C and Shu-Cre, and wild-type counterparts.

Rep1 to 3 indicate each of the replicates (3x 300 eggs each). The observed values (Obs) were calculated as the average of the values of the three replicates. For statistical analysis the observed values were obtained adding up individuals carrying non-recombined and recombined versions of the reporter construct. The expected values (Exp) were calculated as the 25% of the average of the total number of individuals screened in the replicates, considering full survival of the double hemizygous, each of the single hemizygous and wild-type. WT: wild-type. Phenotypes in bold represent the recombined versions.

| | **WT females x Shu-Cre::A-loxN-B-loxN-C_line D males** | | | | |
|---|---|---|---|---|---|
| | **Rep1** | **Rep2** | **Rep3** | **Obs** | **Exp** |
| Shu-Cre::A-loxN-B-loxN-C | 21 | 40 | 30 | 30 | 58 |
| **Shu-Cre::A-loxN-C** | 31 | 41 | 32 | 35 | |
| A-loxN-B-loxN-C | 5 | 2 | 6 | 4 | 58 |
| **A-loxN-C** | 39 | 46 | 38 | 41 | |
| Shu-Cre | 54 | 55 | 55 | 55 | 58 |
| WT | 69 | 69 | 64 | 67 | 58 |
| Total | 219 | 253 | 225 | 232 | 232 |
| | $X^2$= 2.7668, df= 3, p= 0.429 | | | | |

| | **WT females x Shu-Cre::A-loxN-B-loxN-C_line E males** | | | | |
|---|---|---|---|---|---|
| | **Rep1** | **Rep2** | **Rep3** | **Obs** | **Exp** |
| Shu-Cre::A-loxN-B-loxN-C | 18 | 28 | 28 | 25 | 50 |
| **Shu-Cre::A-loxN-C** | 31 | 37 | 33 | 34 | |
| A-loxN-B-loxN-C | 3 | 0 | 1 | 1 | 50 |
| **A-loxN-C** | 51 | 35 | 37 | 41 | |
| Shu-Cre | 35 | 51 | 39 | 42 | 49 |
| WT | 47 | 51 | 69 | 56 | 49 |
| Total | 185 | 202 | 207 | 198 | 198 |
| | $X^2$= 2.2934, df= 3, p= 0.5138 | | | | |

**Supplementary Table 11:** Percentage of individuals of each phenotype observed in the $F_2$ of crosses between double hemizygous of the target lines C or D of A-loxN-B-loxN-C and Shu-Cre, and wild-type counterparts.

Rep1 to 3 indicate each of the replicates (3x 300 eggs each). WT: wild-type. Ave: average percentage amongst the three replicates. S.E.: Standard error. Phenotypes in bold represent the recombined versions.

| | **WT females x Shu-Cre::A-loxN-B-loxN-C_line D males** | | | | |
|---|---|---|---|---|---|
| | **Rep1** | **Rep2** | **Rep3** | **Ave** | **S.E.** |
| Shu-Cre+A-loxN-B-loxN-C | 9.59 | 18.26 | 13.70 | 13.85 | 2.51 |
| **Shu-Cre+A-loxN-C** | 14.16 | 18.72 | 14.61 | **15.83** | 1.45 |
| A-loxN-B-loxN-C | 2.28 | 0.91 | 2.74 | 1.98 | 0.55 |
| **A-loxN-C** | 17.81 | 21.00 | 17.35 | **18.72** | 1.15 |
| Shu-Cre | 24.66 | 25.11 | 25.11 | 24.96 | 0.15 |
| WT | 31.51 | 31.51 | 29.22 | 30.75 | 0.76 |

| | **WT females x Shu-Cre::A-loxN-B-loxN-C_line E males** | | | | |
|---|---|---|---|---|---|
| | **Rep1** | **Rep2** | **Rep3** | **Ave** | **S.E.** |
| Shu-Cre+A-loxN-B-loxN-C | 8.22 | 12.79 | 12.79 | 11.26 | 1.52 |
| **Shu-Cre+A-loxN-C** | 14.16 | 16.89 | 15.07 | **15.37** | 0.81 |
| A-loxN-B-loxN-C | 1.37 | 0.00 | 0.46 | 0.61 | 0.40 |
| **A-loxN-C** | 23.29 | 15.98 | 16.89 | **18.72** | 2.30 |
| Shu-Cre | 15.98 | 23.29 | 17.81 | 19.03 | 2.20 |
| WT | 21.46 | 23.29 | 31.51 | 25.42 | 3.09 |

**Supplementary Table 12:** Sequences of the diagnostic amplicons for the presence/absence of fragment loxN-B-loxN in the experiments analysing the removal of internal sequences.

A-loxN-B-loxN-C: amplicon obtained from the plasmid (positive control), A-loxN-C: amplicon obtained from recombined lines D and E (both sequences were the same, so only one is shown here), purple text: partial sequence of fragment A, black text: *lox*N site, orange text: sequence of fragment B, red text: partial sequence of fragment C, underlined text: sites recognised by the primers.

| | |
|---|---|
| A-loxN-B-loxN-C | **TGTGCAGTCGGTTAGTTGGGAAAGGGCCCTAATTGGGGTAAGTTTTCCCGTTCTTTTCTGGGTTCTTCCCT<br>TTTGCTCATCCTTGCTGCACTACCTTCAGGTGCAAGTCCTAGGATAACTTCGTATAGTATACCTTATACGA<br>AGTTATAACTTAAAAAAAAAAAATCAAAATGGGCAGCCGCCTGGACAAGAGCAAGGTGATCAACAGCGCCC<br>TGGAGCTGCTGAACGAAGTTGGTATCGAGGGCCTGACCACCCGCAAGCTGGCCCAGAAGCTGGGCGTGG<br>AACAGCCGACCCTGTACTGGCACGTGAAGAACAAGCGCGCCCTGCTGGACGCCCTGGCCATCGAAATGC<br>TGGATCGCCACCACACCCACTTCTGCCCGCTGGAGGGCGAGAGCTGGCAGGATTTCCTGCGCAACAACG<br>CCAAGAGCTTCCGCTGCGCCCTGCTGTCGCACCGCGATGGCGCTAAGGTGCACCTGGGCACCCGCCCGA<br>CCGAGAAGCAGTACGAGACCCTGGAGAACCAGCTGGCCTTCCTGTGCCAGCAGGGCTTCAGCCTGGAGA<br>ACGCCCTGTACGCCCTGAGCGCCGTGGGCCACTTCACCCTGGGCTGTGTGCTGGAGGATCAGGAGCACC<br>AGGTGGCCAAGGAGGAGCGCGAGACCCCGACCACCGATAGCATGCCGCCGCTGCTGCGCCAGGCCATCG<br>AGCTGTTCGATCACCAGGGCGCTGAGCCGGCCTTCCTGTTCGGCCTGGAGCTGATCATCTGCGGCCTGGA<br>AAAGCAGCTGAAGTGCGAGAGCGGCAGCGCCTACAGCCGCGCCCGTACCAAGAACAACTATGGCAGCAC<br>CATCGAGGGACTGCTGGACCTGCCGGATGACGATGCCCCGGAGGAAGCCGGCCTGGCCGCCCCCCGCCT<br>GAGCTTCCTGCCCGCCGGACACACGCGCCGCCTGAGCACCGCCCCGCCGACCGACGTTAGCCTGGGCGA<br>CGAGCTGCACCTGGATGGAGAGGATGTGGCAATGGCCCACGCCGACGCCCTGGACGATTTCGACCTGGA<br>TATGCTGGGCGATGGAGATAGCCCGGGACCGGGCTTCACGCCCCACGATAGCGCCCCGTACGGCGCTCT<br>GGACATGGCCGACTTCGAGTTCGAGCAAATGTTCACCGACGCGCTGGGCATCGATGAGTACGGCGGTTA<br>ATTAATTGTTAAGATACATTGATGAGTTTGGACAAACCACAACTAGAATGCAGTGAAAAAAATGCTTTATT<br>TGTGAAATTTGTGATGCTATTGCTTTATTTGTAACCATTATAAGCTGCAATAAACAAGTTAACAACAACAA<br>TTGCATTCATTTTATGTTTCAGGTTCAGGGGGAGGTGTGGGAGGTTTTTTAAAGCAAGTAAAACCTCTACA<br>AATGTGGTATGGCTGATTATGATCAGGCCAGGGCGCTGGGGAAGGCGATGGCGTGCTCGGTCAGCTGCC<br>ACTTCTGGTTCTTGGCGTCGCTCCGGTCCTCCCGCAGCAGCTTGTGCTGGATGAAGTGCCACTCGGGCAT<br>CTTGCTGGGCACGCTCTTGGCCTTGTACACGGTGTCGAACTGGCACCGGTACCGGCCGCCGTCCTTCAGC** |

| | |
|---|---|
| | **AGCAGGTACATGCTGACATCGCCCTTCAGGATGCCCTGCTTAGGCACGGGCATGATCTTCTCGCAGCTGG<br>CCTCCCAGTTGGTGGTCATCTTCTTCATCACGGGGCCGTCGGCGGGGAAGTTCACGCCGTTGAAGATGCT<br>CTTGTGGTAGATGCAGTTCTCCTTCACGCTCACGGTGATGTCCACGTTACAGATGCACACGGCGCCGTCC<br>TCGAACAGGAAGCTCCGGCCCCAGGTGTAGCCGGCGGGGCAGCTGTTCTTGAAGTAGTCCACGATGTCC<br>TGGGGGTACTCGGTGAAGATCCGGTCGCCGTACTTGAAGCCGGCGCTCAGGATGTCCTCGCTGAAGGGC<br>AGGGGGCCGCCCTCGATCACGCACAGGTTGATGGTCTGCTTGCCCTTGAAGGGGTAGCCGATGCCCTCG<br>CCGGTGATCACGAACTTGTGGCCGTTCACGCAGCCCTCCATGTGGTACTTCATGGTCATCTCCTCCTTCA<br>GGCCGTGCTTGCTGTGGGCCATTCGACTCTAGCGGTACCCCGATTGTTTAGCTTGTTCAGCTGCGCTTGT<br>TTATTTGCTTAGCTTTCGCTTAGCGACGTGTTCACTTTGCTTGTTTGAATTGAATTGTCGCTCCGTAGACG<br>AAGCGCCTCTATTTATACTCCGGCGGTCGAGGGTTCGAAATCGATAAGCTTGGATCCTAATTGAATTAGC<br>TCTAATTGAATTAGTCTCTAATTGAATTAGATCCCCGGGCCCAGATCTTTCGACTTTCACTTTTCTCTATCA<br>CTGATAGGGAGTGGTAAACTCGACTTTCACTTTTCTCTATCACTGATAGGGAGTGGTAAACTCGACTTTCA<br>CTTTTCTCTATCACTGATAGGGAGTGGTAAACTCGACTTTCACTTTTCTCTATCACTGATAGGGAGTGGTA<br>AACTCGACTTTCACTTTTCTCTATCACTGATAGGGAGTGGTAAACTCGACTTTCACTTTTCTCTATCACTG<br>ATAGGGAGTGGTAAACTCGACTTTCACTTTTCTCTATCACTGATAGGGAGTGGTAAACTCGAAAACGAGC<br>GCCGGAGTATAAATAGAGGCGCTTCGTCTACGGAGCGACAATTCAATTCAAACAAGCAAAGTGAACACGT<br>CGCTAAGCGAAAGCTAAGCAAATAAACAAGCGCAGCTGAACAAGCTAAACAATCTGCGGTACCCTGGCGT<br>CTAATTGGGGTAAGTTTTCCCGTTCTTTTCTGGGTTCTTCCCTTTTGCTCATCCTTGCTGCACTACCTTCA<br>GGTGCAAGTAAAGCCCGCGGTGGGAGGCCGGCCGATAACTTCGTATAGTATACCTTATACGAAGTTATCG<br>CCGCCACCATGCCCACCCCACCCAAGAAGAAGCGCAAAGCTAGCGTTATGGCCCTGTCCAACAAGTTCAT<br>CGGCGACGACATGAAGATGACCTACCACATGGACGGCTGCGTGAACGGCCACTACTTCACCGTGAAGGG<br>CGAGGGCAGCGGCAAGCCCTACGAGGGCACCCAGACCTCCACCTTCAAAGTCACAATGGCC** |
| A-loxN-C | **TGTGCAGTCGGTTAGTTGGGAAAGGGCCCTAATTGGGGTAAGTTTTCCCGTTCTTTTCTGGGTTCTTCCCT<br>TTTGCTCATCCTTGCTGCACTACCTTCAGGTGCAAGTCCTAGGATAACTTCGTATAGTATACCTTATACGA<br>AGTTATCGCCGCCACCATGCCCACCCCACCCAAGAAGAAGCGCAAAGCTAGCGTTATGGCCCTGTCCAAC<br>AAGTTCATCGGCGACGACATGAAGATGACCTACCACATGGACGGCTGCGTGAACGGCCACTACTTCACCG<br>TGAAGGGCGAGGGCAGCGGCAAGCCCTACGAGGGCACCCAGACCTCCACCTTCAAAGTCACAATGGCC** |

**Supplementary Table 13:** Sequences of the *attP* and *attB* sites and the diagnostic amplicons for the confirmation of PhiC31 mediated integration.

Blue text: sequence of the *attP* site, red text: sequence of the *attB* site, orange text: partial sequence of the target site, black text: partial sequence of the inserted transgene, underlined text: sites recognised by the primers, text in italics: sections of *attP* and *attB* that form *attL*.

| | |
|---|---|
| *attP* | ***AGTAGTGCCCCAACTGGGGTAACCTTTG*****AGTTCTCTCAGTTGGGGGCGTA** |
| *attB* | **TCGACGATGTAGGTCACGGTCTCGAAGCCGCGGTGCGGGTGCCAGGGCGTGCCCTTG*****GGCTCCCCGGGCGCGTACT**<br>***CCACCTCACCCATCTGGTCCATCATGATGAACGGGTCGAGGTGGCGGTAGTTGATCCCGGCGAACGCGCGGCGCACCGGGA***<br>***AGCCCTCGCCCTCGAAACCGCTGGGCGCGGTGGTCACGGTGAGCACGGGACGTGCGACGGCGTCGGCGGGTGCGGATACGC***<br>***GGGGCAGCGTCAGCGGGTTCTCGACGGTCACGGCGGGCATGTCGAC*** |
| *attL* | **TCGGTCTGTATATCGAGGTTTATTTATTAATTTGAATAGATATTAAGTTTTATTATATTTACACTTACATACTAATAA**<br>**TAAATTCAACAAACAATTTATTTATGTTTATTTATTTATTAAAAAAAAACAAAAACTCAAAATTTCTTCTATAAAGTA**<br>**ACAAAACTTTTATCGAATTGCTTCGGCGCCA*****AGTAGTGCCCCAACTGGGGTAACCTTTGGGCTCCCCGGGCGCGTACTCC***<br>***ACCTCACCCATCTGGTCCATCATGATGAACGGGTCGAGGTGGCGGTAGTTGATCCCGGCGAACGCGCGGCGCACCGGGAAG***<br>***CCCTCGCCCTCGAAACCGCTGGGCGCGGTGGTCACGGTGAGCACGGGACGTGCGACGGCGTCGGCGGGTGCGGATACGCG***<br>***GGGCAGCGTCAGCGGGTTCTCGACGGTCACGGCGGGCATGTCGAC*****TCTAGGCGCGCCTCGGATTCGACTTTCACTTTTC**<br>**TCTATCACTGATAGGGAGTGGTAAACTCGACTTTCACTTTTCTCTATCACTGATAGGGAGTGGTAAACTCGACTTTC**<br>**ACTTTTCTCTATCACTGATAGGGAGTGGTAAACTCGACTTTCACTTTTCTCTATCACTGATAGGGAGTGGTAAACTC**<br>**GACTTTCACTTTTCTCTATCACTGATAGGGAGTGGTAAACTCGACTTTCACTTTTCTCTATCACTGATAGGGAGTGG**<br>**TAAACTCGACTTTCACTTTTCTCTATCACTGATAGGGAGTGGTAAACTCGAAAACGAGCGCCGGAGTATAAATAGA**<br>**GGCGCTTCGTCTACGGAGCGACAATTCAATTCAAACAAGCAAAGTGAACACGTCGCTAAGCGAAAGCTAAGCAAAT**<br>**AAACAAGCGCAGCTGAACAAGCTAAACAATCTGCGGTACCCTGGCGTCTAATTGGGGTAAGTTTTCCCGTTCTTTT**<br>**CTGGGTTCTTCCCTTTTGCTCATCCTTGCTGCACTACCTTCAGGTGCAAGTAAAGCCATACAGTGAACCAGGCCACC**<br>**ATGGGAGATCCCACCCCACCCAAGAAGAAGCGCAAAGCTAGCGTTATGGCCCTGTCCAACAAGTTCATCGGCGACG**<br>**ACATGAAGATGACCTACCACATGGACGGCTGCGTGAACGGCCACTACTTCACCGTGAAGGG** |
| *attR* | **GCACAAGCTGGAGTACAACTACATCAGCCACAACGTCTATATCACCGCCGACAAGCAGAAGAACGGCATCAAGGCC**<br>**AACTTCAAGATCCGCCACAACATCGAGGACGGCAGCGTGCAGCTCGCCGACCACTACCAGCAGAACACCCCCATCG**<br>**GCGACGGCCCCGTGCTGCTGCCCGACAACCACTACCTGAGCACCCAGTCCGCCCTGAGCAAAGACCCCAACGAGA**<br>**AGCGCGATCACATGGTCCTGCTGGAGTTCGTGACCGCCGCCGGGATCACTCTCGGCATGGACGAGCTGTACAAGT**<br>**AAAGCGGCCGCGACTCTAGATAACTGGAGCTTGATAACATTATACCTAAACCCATGGTCAAGAGTAAACATTTCTG**<br>**CCTTTGAAGTTGAGAACACAATTAAGCATCCCCTGGTTAAACCTGACATTCATACTTGTTAATAGCGCCATAAACAT** |

**AGCACCAATTTCGAAGAAATCAGTTAAAAGCAATTAGCAATTAGCAATTAGCAATAACTCTGCTGACTTCAAAACGA
GAAGAGTTGCAAGTATTTGTAAGGCACAGTTTATAGACCACCGACGGCTCATTAGGGCTCGTCATGTAACTAAGCG
CGGTGAAACCCAATTGAACATATAGTGGAATTATTATTATCAATGGGGAAGATTTAACCCTCAGGTAGCAAAGTAAT
TTAATTGCAAATAGAGAGTCCTAAGACTAAATAATATATTTAAAAATCTGGCCCTTTGACCTTGCTTGTCAGGTGCA
TTTGGGTTCAATCGTAAGTTGCTTCTATATAAACACTTTCCCCATCCCCGCAATAATGAAGAATACCGCAGAATAAA
GAGAGATTTGCAACAAAAAATAAAGGCATTGCGAAAACTTTTTATGGGGGATCATTACACTCGGGCCTACGGTTAC
AATTCCCAGCCACTTAAGCGACAAGTTTGGCCAACAATCCATCTAATAGCTAATAGCGCAATCACTGGTAATCGCAA
GAGTATATAGGCAATAGAACCCATGGATTTGACCAAAGGTAACCGAGACAATGGAGAAGCAAGAGGATTTCAAACT
GAACACCCACAGTACTGTGTACTACCACTGGCGCGTTTGGGACCTGCAGATAACTTCGTATAGTATACCTTATACG
AAGTTATCGATTGTTTCCTAGGTCGACGATGTAGGTCACGGTCTCGAAGCCGCGGTGCGGGTGCCAGGGCGTGCC
CTTGAGTTCTCTCAGTTGGGGGCGTAGGTCGACAAGCTTTACGAGTAGAATTCTACGCGTAAAACACAATCAAGTA
TGAGTCATAATCTGATGTCATGTTTTGTACACGGCTCATAACCGAACTGG**
